# Master regulators of genetic interaction networks mediating statin drug response in *Saccharomyces cerevisiae* vary with genetic background

**DOI:** 10.1101/443879

**Authors:** Bede P. Busby, Eliatan Niktab, Christina A. Roberts, Namal V. Coorey, Jeffrey P. Sheridan, Dinindu S. Senanayake, Andrew B. Munkacsi, Paul H. Atkinson

## Abstract

Determination of genetic interaction networks (GINs) surrounding drug targets identifies buffering genes and provides molecular insight into drug response in individuals. Here we used backcross methodology to create *Saccharomyces cerevisiae* deletion libraries in three genetic backgrounds resistant to statins, which are additional to the statin-sensitive S288C deletion library that has provided much of what is known about GINs in eukaryotes. Whole genome sequencing and linkage group analysis confirmed the genomic authenticity of the new deletion libraries. Statin response was probed by drug-gene interactions with atorvastatin and cerivastatin treatments, as well as gene-gene interactions with the statin target *HMG1* and *HMG2* genes or the sterol homeostatic *ARV1* gene. The 20 GINs generated from these interactions were not conserved by function or topology across the four genetic backgrounds. Centrality measures and hierarchical agglomerative clustering identified master regulators that if removed collapsed the networks. Community structure distinguished a characteristic early secretory pathway pattern of gene usage in each genetic background. ER stress in statin-resistant backgrounds was buffered by protein folding genes, which was confirmed by reduced activation of the unfolded protein response in statin-resistant backgrounds relative to the statin-sensitive S288C background. These network analyses of new gene deletion libraries provide insight into the complexity of GINs underlying individual drug response.

## INTRODUCTION

Understanding phenotypes that are genetically complex requires analysis of multiple genes contributing to phenotypes. Classical genetic studies have long investigated the additive functions of genes in polygenic or quantitative traits ^1,2^, the effects of modifying alleles ^3,4^, genetic capacitators ^5^, “missing heritability” ^6^, the epistatic contribution to phenotype ^7,8^, as well as both additive and epistatic effects ^9^. In an advance using systems biology, gene functions have also been implied in genetic interaction networks (GINs) comprising overlapping synthetic lethal epistatic interactions (GIs) between pairs of non-essential gene deletions identified in high throughput by synthetic genetic array (SGA) technology ^10^. Epistasis occurs because some gene pairs have functional commonality and, logically, specific GINs of overlapping epistatic gene pairs can also be considered functional ^11–13^. We thus raise the question: Are there specific functional GINs that are conserved or do they vary in individuals?

The debate is open as the conservation and effects of perturbation of these GINs remains largely predictive ^14^ via comparisons of specific synthetic lethal GIs in human and yeast ^15^, GIs in the distantly related yeast and *Caenorhabditis elegans* ^16,17^, essential genes in two strains of *S. cerevisiae* ^4^, and protein-protein interactions in *S. pombe* and *S. cerevisia*e ^18^. There is broad concurrence that specific genes of GINs are not well conserved but that some GIs and network topological features are conserved ^11^. However, systematic analyses of GINs have not been investigated in multiple genetic backgrounds of the same yeast species utilising deletion libraries specific for each genetic background and we address this question here for statin-specific GINs.

Statins target HMG-CoA-reductase and are amongst the most prescribed of all therapeutic drugs ^19^. However, they have significant side-effects in some individuals such as muscular myopathies ^20,21^ and exhibit individual variation in clinical efficacy ^22^. Atorvastatin and cerivastatin are two cholesterol-lowering drugs that have the same target, HMG-CoA reductase (encoded by *HMGCR* in humans and the orthologous paralogues *HMG1* and *HMG2* in yeast), but cerivastatin is no longer FDA-approved owing to adverse side effects, suggesting that analysis of GINs in response to these drugs in different individual genetic backgrounds could be critical to fully understanding the mechanisms of these drugs. The statin drug target and sterol pathways are conserved from yeast to humans ^23^, the *HMG1/2* deletion has been used as a genetic mimic of statin treatment ^24,25^, chemical genomic analyses in yeast elucidated the cellular response to statins ^26^, and genome-wide analyses in yeast identified GIs with the *HMG1/2* statin drug target ^13^.

Deletion libraries in the yeast S288C background used in synthetic genetic array (SGA) and chemical genetic analyses have provided much of what is known about GINs in eukaryotes ^10,13,27,28^. In this paper we extended the scope of the S288C literature by creating three new deletion libraries in three additional yeast strains of different genetic backgrounds. We identified GIs with atorvastatin, cerivastatin, and statin target *HMG1*, its functional paralog *HMG2*, and the sterol homeostatic *ARV1* genes as query genes in SGAs in four genetic backgrounds that generated 20 GINs. We found that individual strains had highly variable GINs by multiple criteria including: functional, topological, and clustering comparisons. We found that statins, in resistant backgrounds, were buffered by protein folding genes, which was confirmed by observation of reduced activation of the unfolded protein response relative to the statin-sensitive S288C background. Our results provide experimental and computational resources that model the complexity of GINs underlying individual drug response.

## RESULTS

### Selection of statin-resistant strains

We obtained 36 haploid derivatives (Supplementary Table S1) from the fully-sequenced wild-type strains in the Saccharomyces Genome Resequencing Project (SGRP) collection ^29^ and evaluated growth via spot dilutions on agar plates containing increasing concentrations of atorvastatin or cerivastatin. There was a large range of phenotypic diversity observed for these strains with some being ^~^10-fold more sensitive and others ^~^10-fold more resistant relative to BY4742 (the wild-type strain in the S288C background). Three strains (Y55, UWOPS87-2421 and YPS606) exhibited normal growth at 400 μM atorvastatin in contrast to 50 μM atorvastatin that was lethal to S288C (Figure 1A). Likewise, growth of these strains at 80 μM cerivastatin was comparable with growth of S288C at 50 μM cerivastatin (Figure 1B). These strains were chosen for further investigation.

**Figure 1.**
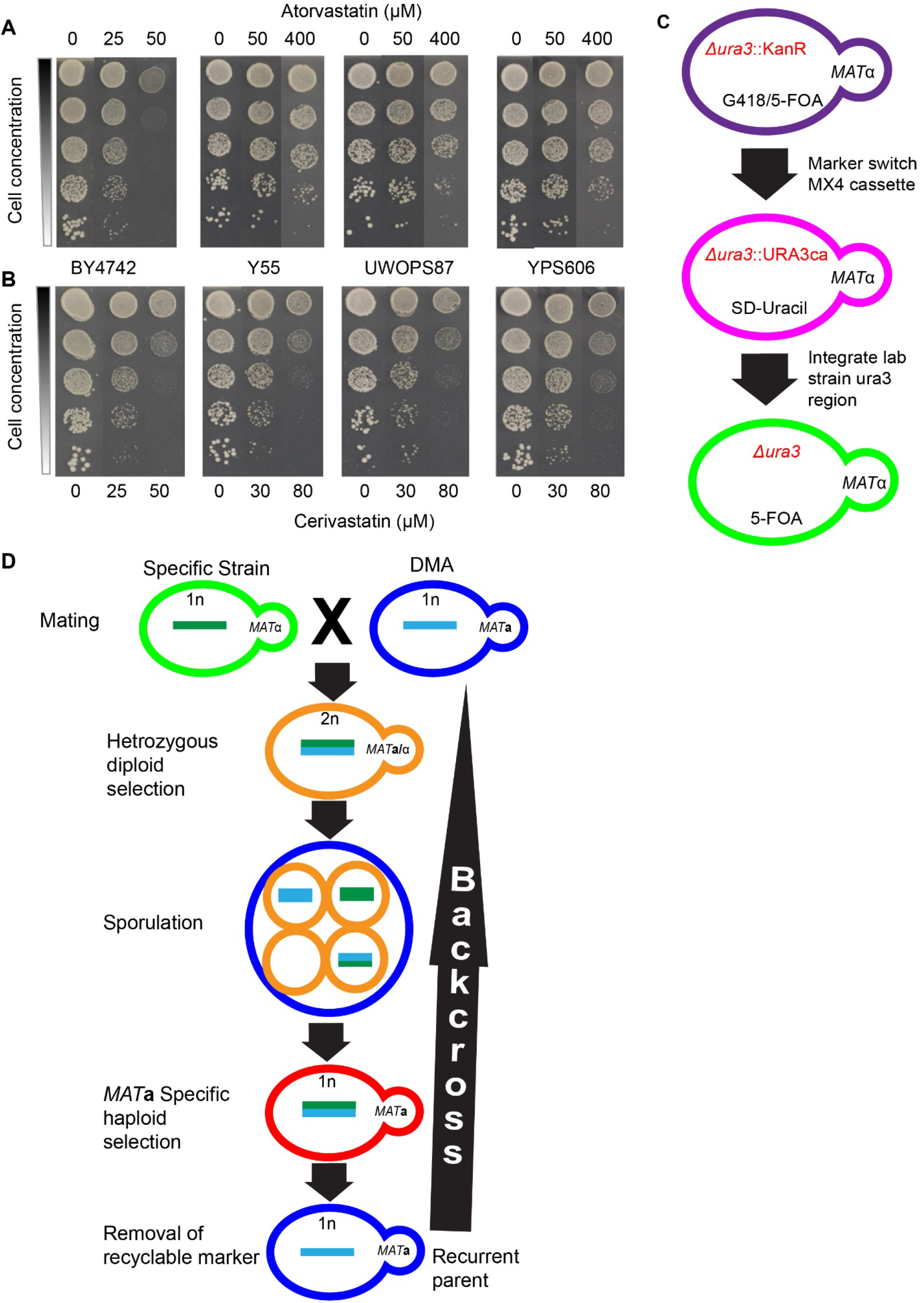
Statin resistance of SGRP strains and backcross methodology to construct ssDMA libraries that represent genome-wide deletion libraries of these SGRP strains. A-B) Serial dilutions of three SGRP strains (Y55, UWOPS87-2421 and YPS606) resistant to A) atorvastatin and B) cerivastatin relative to BY4742 in the S288C background. C) Outline of marker switch method used to introduce the *URA3* marker into the SGRP strains. D) Outline of method used to cross and backcross the SGRP strains with the S288C deletion library (DMA) to produce the ssDMA libraries.

### Construction and validation of three new strain deletion mutant arrays

To investigate GINs underlying resistance of these strains to statins, we constructed new strain-specific genome-wide deletion libraries for the three statin-resistant strains (Y55, UWOPS87-2421 and YPS606) for subsequent use in chemical genetic and SGA analyses. First, we used a marker-switching strategy to introduce *ura3-Δ* and *his3-Δ* into the SGRP strains (Figure 1C, Supplementary Table S1). Then we used selection markers within the SGA methodology ^28^ to cross and backcross these strains six times with the S288C deletion library (Figure 1D). The SGRP strains for making these libraries had sporulation rates of 70-80% compared to only 3% in S288C, hence greatly facilitating the sporulation step in SGAs with the new deletion libraries. We named the new deletion libraries <u>S</u>RGP <u>s</u>train <u>d</u>eletion <u>m</u>utant <u>a</u>rrays (ssDMAs), which retained the statin-resistant phenotype (Supplementary Figure S1).

Whole-genome sequencing of the new strains was performed from 384 pooled colonies of a specific plate (plate #10) from each of the three ssDMAs and the S288C deletion library, which were further compared with the S288C haploid derivative BY4742. Each ssDMA sequence had a 0.2-0.4% higher alignment score to its SGRP parental strain than to the BY4742 parental strain (Supplementary Table S2). Furthermore, each ssDMA sequence had more regions with perfect (100%) alignment with its UWOPS87-2421, Y55 and YPS606 SGRP parent strains than the S288C parent (Supplementary Figures S2-S4) confirming that the background of each ssDMA closely resembled its SGRP parent. Synteny analysis showed that there were not any major structural rearrangements in the ssDMA or parental SGRP strains (Supplementary Figures S5-S7) Synteny with S288C was also supported with the recovery of the expected linkage groups surrounding the three query genes in the SGAs later described (Supplementary Table S3). Analysis of the sequencing coverage indicated that each ssDMA was euploid, consistent with the controls (Supplementary Figure S8). Genetic variation between the strains was also evident revealed by 5.9, 5.4 and 5.9 SNPs/kb in Y55, UWOPS87-2421 and YPS606, respectively, a finding that generally exceeded the SNP density among ethnically distinct human populations ^30^, thus indicating there is extensive genetic diversity in our yeast backgrounds around which to investigate genetic background effects on GINs.

### Chemical genetic and genetic interaction profiles differ between yeast strain strains

The central tenet of chemical genetics ^31^ is that a drug or small molecule inhibitor (SMI) binds specifically to a gene product and alters or ablates its function mimicking a mutation. Thus, an SMI can be paired with a deletion mutation to screen for hypersensitive epistatic interaction profiles *en masse* in deletion libraries to provide information on the target and buffering mechanisms of the SMI ^27^. We obtained chemical genetic profiles of our four deletion libraries via growth measurements in the presence and absence of statin treatment (25 μM atorvastatin or 10 μM cerivastatin for the S288C DMA, 100 μM atorvastatin or 50 μM cerivastatin for the statin-resistant ssDMAs). These concentrations inhibited the susceptible and resistant strain growth to approximately the same amount (Figure 1A). The chemical genetic profiles for atorvastatin (Figure 2A) and cerivastatin (Figure 2B) were substantially different in the four strains. In the profile of 288 gene deletions hypersensitive to atorvastatin, only four deletions (*HMG1, MID1, MID2, PDR1*) were common to all four strains. In the profile of 283 gene deletions hypersensitive to cerivastatin, only three gene deletions (*HMG1, HMS2, SPF1*) were common in all four strains. The overall profiles were distinct in the different strains because most of the hypersensitive gene deletions were unique to any one strain (Supplementary Table S4). Only one interaction for atorvastatin was common to the three resistant strains and not S288C; this was *SLT2*, a MAP kinase regulator of proteasome abundance that has GIs with the major protein folding sensors *IRE1* and *HAC1* ^32^ as well as with *CWH41*, a glycosylation processing glucosidase critical to sensing protein folding in the ER ^33^.

**Figure 2.**
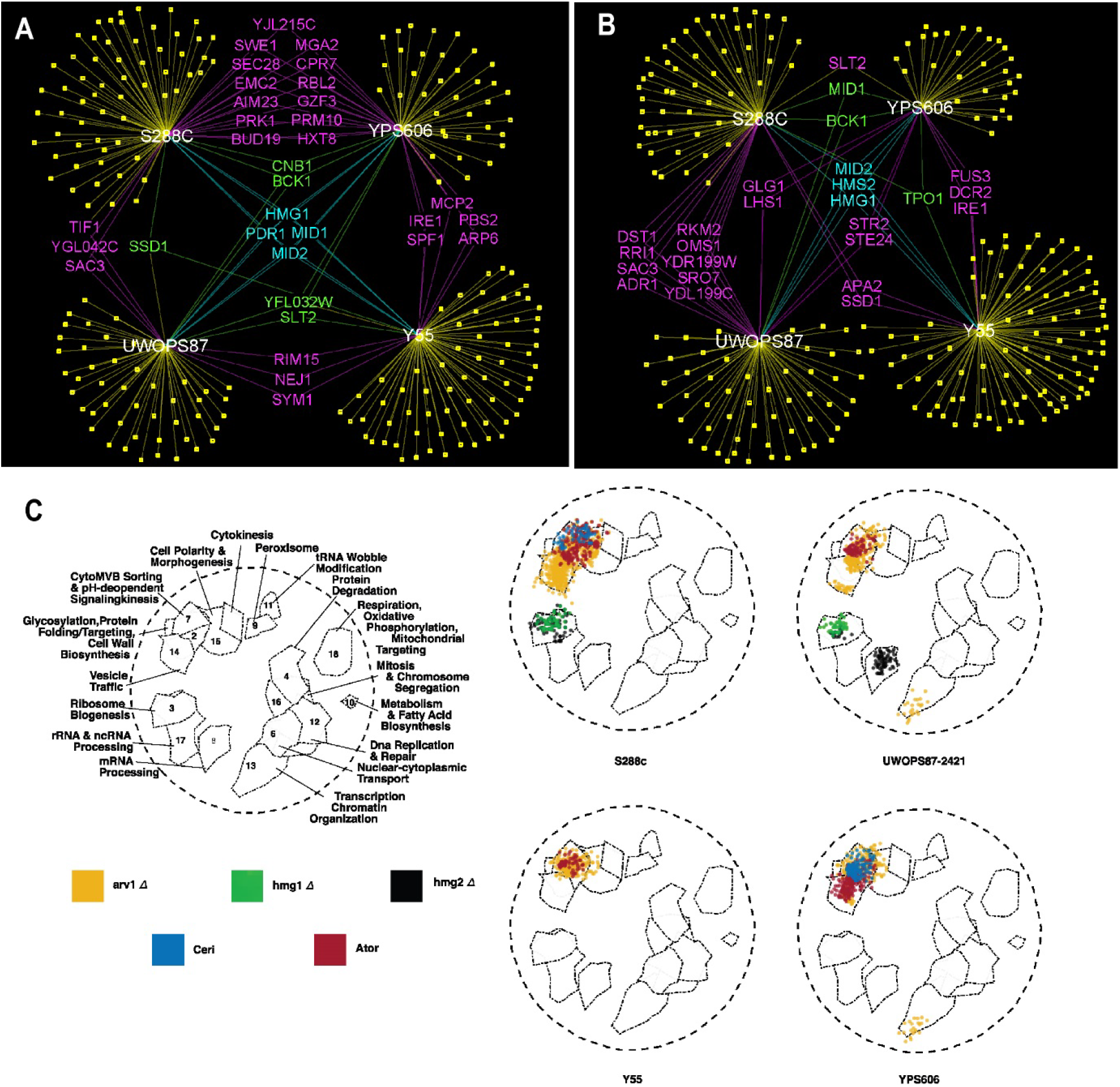
Primary genetic interaction networks and functional annotation illustrates variation across genetic backgrounds. A-B) Chemical genetic profile for A) atorvastatin and B) cerivastatin representing interactions shared by different genetic backgrounds (purple, green, and blue) or interactions unique to specific genetic backgrounds (yellow, genes not included in this figure). C) Spatial Analysis of Functional Enrichment (SAFE) of 20 GINs derived from five queries (atorvastatin, cerivastatin, *hmg1*-*Δ*, *hmg2-Δ*, *arv1-Δ*) and four genetic backgrounds (S288C, UWOPS87-2421, Y55, YPS606). Relative to the 18 functional regions previously defined^13^, the figure illustrates significant membership of each GIN to these functional regions where the intensity of the colour is proportional to the number of interactions.

Genes involved in the unfolded protein response (UPR) were required in the response to atorvastatin in the three resistant strains (Figure 2A). In yeast, the major mediator of UPR is *IRE1*, which upon ER unfolded protein stress, is released from the Hsp70 chaperone *KAR2* in the ER whence it oligomerises and activates the UPR pathway ^34^. We found the *ire1-Δ* strain was hypersensitive to atorvastatin in Y55 and YPS606, while it was hypersensitive to cerivastatin only in Y55 (Figure 2A-B). The nucleotide exchange factors for additional UPR genes *KAR2*, *SIL1* and *LHS1* ^35^ were also required in the resistant strains. Specifically, *sil1-Δ* was hypersensitive to atorvastatin in UWOPS87-2421 and *lhs1-Δ* was hypersensitive to cerivastatin in UWOPS87-2421 and YPS606 (Figure 2A). Notably, neither the deletion of *IRE1* nor *SIL1* conferred hypersensitivity to atorvastatin or cerivastatin in the susceptible S288C strain (Supplementary Table S4). These several UPR genes show synthetic lethality as double deletions such as *ire1-Δ/lhs1-Δ*, *ire1-Δ/sil1-Δ* and *lhs1-Δ/sil1-Δ* ^36^, emphasizing their interdependent relationship to ER folding processes.

We created strain-specific GINs utilising the query strains *hmg1-Δ*, *hmg2-Δ* and *arv1-Δ*, since all three genes are fundamental to sterol homeostasis in the ER, synthetic lethal with *IRE1* ^36–38^ or involved with glycosylphosphatidylinositol (GPI) biosynthesis, which is one of major branches of the mevalonate pathway. The 12 SGA procedures generated 50,400 unique double deletion mutants, of which 37,800 have not been previously constructed. The negative GIs (synthetic lethal and synthetic sick interactions) for each background were scored as double deletion strains (*Z*-score > 2.0, *P*-value < 0.05) with significant growth defects in the double mutants relative to the single deletion mutants. Genes in the linkage groups of the *hmg1-Δ*, *hmg2-Δ* and *arv1-Δ* queries were not included as GIs (Supplementary Table S3). Across the four genetic background, 339 GIs were identified with the *hmg1-Δ* query, 181 GIs with the *hmg2-Δ* query, and 425 GIs with the *arv1-Δ* query (Supplementary Table S4; Supplementary Figures S9-S11). There was only limited overlap in the four genetic backgrounds of GIs for particular query strains.

Specifically, the *hmg1-Δ* query gene exhibited overlapping GIs with *hmg2-Δ* in all backgrounds (Supplementary Figure S9), GIs unique to the resistant strains (*e.g.*, *SEC66, HTZ1, NUM1, YML102C, SHE4* and *IRE1*) were common to two resistant strains Y55 and UWOPS87-2421 and GIs shared by sensitive and resistant backgrounds (*SAC1, SPF1* and *GIM5*) were common to the sensitive S288C as well as the resistant Y55 and UWOPS87-2421. For the *hmg2-Δ* query, only *HMG1* was a common GI to all four strains (Supplementary Figure S10) and the helicase *YRF1-6* was common to two resistant strains (Y55, UWOPS87-2421). The *arv1-Δ* query gene exhibited GIs only with *SLT2, CDC73, CHS3, CHS7*, and *IRE1* in all four strains and GIs shared by only the resistant strains were with *LEO1, YBL062W, RPS8A, SKT5* and *HMG1* (Supplementary Figure S11). There was a requirement for *SPF1* in the resistant strains YPS606 and Y55 in the presence of atorvastatin and cerivastatin, but not in S288C. Interestingly, *SPF1* is an ER ATPase ion transporter that regulates *HMG2* degradation ^39^ and interacts with the UPR regulators *HAC1* and *IRE1* ^32^, implicating buffering by the ER UPR system in the statin drug response. The various GIs arising from the statin chemical genetic profiles and the SGA queries indicated, among other processes, a clear involvement of early secretory pathway processes which we further investigated in this paper.

For functional and topological analyses we augmented our 20 primary GINs (Supplementary Table S4) utilising the single most comprehensive GIN for any eukaryote ^13^ which is a systematic, genome-wide yeast GIN comprising ~23 million GIs to integrate the 1660 GIs of our 20 primary GINs representing the five queries (atorvastatin, cerivastatin, *hmg1-Δ, hmg2-Δ, arv1-Δ*) in the four genetic backgrounds. Using adjacency matrices ^40^, we added all genes in path-lengths of 3 emanating from the hit vertices of the primary GINs (Supplementary Table S4) by utilising *K*-edge centralities. The resulting 20 adjacency matrix files of ^~^2000 x ^~^2000 nodes each made up our 20 augmented GINs (Supplementary Table S5) were used for all analyses in the rest of this paper.

### SAFE analysis reveals functional heterogeneity of gene usage among yeast strains

Using the 20 augmented GINs, we performed Spatial Analysis of Functional Enrichment (SAFE) analysis ^12,13^ (Supplementary Table S6) to elucidate functional properties of the GINs. SAFE is a Gene Ontology (GO) functional depiction analysis with network topology constrained graph distance similarities that represents statistically and quantitatively localised enrichments of genes in specific cellular processes ^12^ (Figure 2C). The atorvastatin treatment shows a concentration of GIs enriched in the early secretory pathway related Region 2 (*glycosylation/protein folding*) and Region 7 *(multi vesicular body (MVB) sorting*) in all four strains. Notably different to the other strains, the resistant strain YPS606 was also significantly enriched in Region 14 (*vesicle trafficking*). In contrast, the cerivastatin treatment showed a concentration of GIs enriched in Region 2 (*glycosylation/protein folding*) and Region 7 (*MVB sorting*) only in S288C and Y55, and Region 15 (*cell polarity and morphogenesis*) only in S288C. These results show that not only do the strains differ functionally but that the cerivastatin response is markedly different to atorvastatin.

Consistent with the SAFE chemical genetic profiles, SAFE analyses of GIs with the query genes *HMG1, HMG2* and *ARV1* also showed extensive heterogeneity of gene usage across the four genetic backgrounds. The GIs with *ARV1* were enriched only in Region 7 (*MVB sorting*) in all four strains, while there was enrichment in Region 2 (*glycosylation/protein folding*) and Region 14 (*vesicle trafficking*) in two resistant strains (YPS606 and UWOPS87-2421) whilst the statin-sensitive S288C GIs were concentrated in Region 14 (*vesicle trafficking*). Unique to YPS606 was an enrichment in Region 13 (*transcription, chromatin organization).* The GIs with *HMG1* and *HMG2* were enriched in Region 3 (*ribosome biogenesis*) in only two strains (S288C and UWOPS87-2421) with a strikingly unique signature in Region 8 (*mRNA processing*) in the UWOPS87-2421 strain.

### Hierarchical functional clustering shows mevalonate pathway varies between strains

We next elucidated functional similarities among chemical genetic and GI profiles by employing agglomerative and *k*-means hierarchical clustering ^41^. We distinguished three major clades (Figure 3A). The first clade had five GI profiles; these were GI profiles only in the S288C and YPS606 strains (GIs with *HMG1* and *HMG2* in YPS606 as well as GIs with *HMG1*, *HMG2* and *ARV1* in S288C). The second clade comprised six chemical genetic profiles (particularly the atorvastatin chemical genetic profile in all four backgrounds but only the cerivastatin chemical genetic profile in UWOPS87-2421 and YPS606) and six GI profiles (GIs with *ARV1* in Y55 and YPS606, GIs with *HMG1* in Y55 and UWOPS87-2421, GIs with *HMG2* in Y55 and UWOPS87-2421). The third clade comprised two chemical genetic profiles (cerivastatin in S288C and Y55) and the GIs with *ARV1* in UWOPS87-2421. These results indicate that S288C is functionally more closely related to YPS606 and Y55 is more closely related to UWOPS87-2421, while Y55 and UWOPS87-2421 were markedly different.

**Figure 3.**
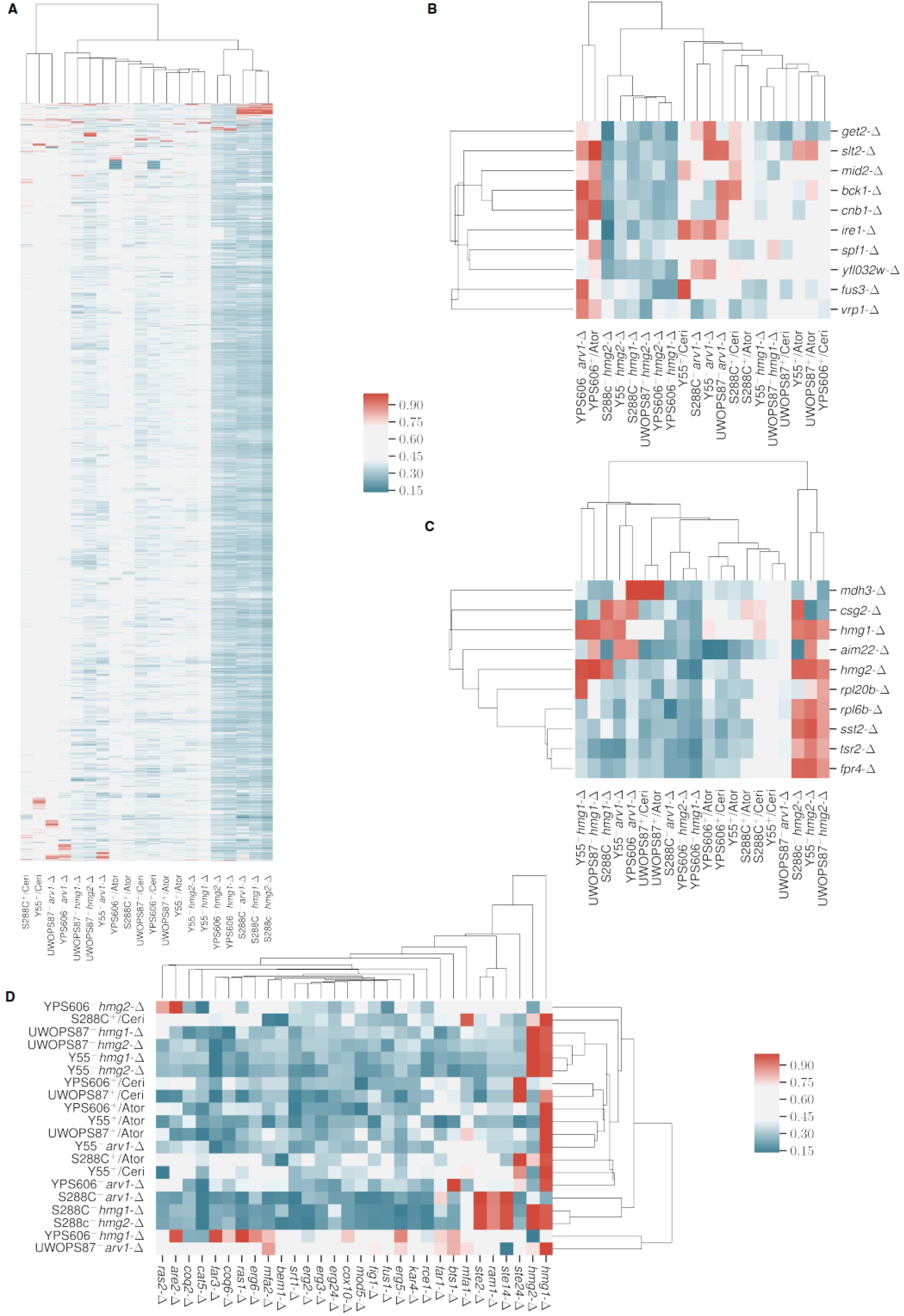
Hierarchical clustering of GINs represents overall genetic background variation throughout the genome and specifically the unfolded protein response and mevalonate pathway. A-D) Hierarchical clustering of growth phenotypes representing A) genome-wide deletion libraries, B) *IRE1* (the major mediator of UPR), C) *HMG1* and *HMG2* (the targets of atorvastatin and cerivastatin), and D) genes within and branching from the mevalonate pathway. One scale applies to A-C and the scale for D shown separately to the right of the cluster. All scales represent min-max normalised *Z*-scores.

We narrowed the focus of Figure 3A to 10 contiguous genes re-clustering around *IRE1* the major mediator of UPR. Consistent with clustering of the whole genome, there was clear evidence of strain-dependent clustering of genes such as clusters focussed around *IRE1* (Figure 3B) from the *hmg2-∆* query in S288C, Y55 and UWOPS87-2421 strains that were distinct from the *hmg2-∆* query in YPS606 (Figure 3B). *IRE1* clustered closely with *SPF1* across all backgrounds (Figure 3B), reiterating the primordial association of sterol/lipid homeostasis and the UPR. Since *SPF1* is synthetic lethal with *BTS1* ^32^, a major branch point in the mevalonate pathway and also has GIs with many genes involved in UPR (*e.g.*, *HAC1, HMG1, IRE1*, *ARV1*, many *COG* genes, *CWH41*, numerous ERG genes and all three *GET* genes)^32^, we suggest that *SPF1* is a functional hub interacting with many genes of the ER folding pathways, the mevalonate pathway and the early secretory pathway. In a second focussed re-clustering around the statin target *HMG1*, strain-dependency was again detected. *HMG1* clustered strongly with *CSG2* (an ER protein involved in sphingolipid synthesis that has GI with *HMG1*) and *AIM22* (a mediator of mitochondrial inheritance and protein lipoylation) only in Y55 and UWOPS87-2421 (Figure 3C).

The third refocussed cluster analysis was based on relationship to a panel of genes we curated from the mevalonate pathway and three main branches namely the ergosterol branch, dolichol N-glycan synthesis branch, and the all-trans GGPP isoprenoid branch of protein prenylation (Supplementary Table S4). In these clusters (Figure 3D), one clade contains GIs with *ARV1* in the background of UWOPS87-2421 and *HMG1* in the background of YPS606, similarly to that seen in Figure 3A. For the query genes *hmg1-∆, hmg2-∆* and *arv1-∆* in the S288C genetic background, there is a cluster containing strong negative GIs with *STE14* (ER protein mediating processing of alpha-factor and RAS proteins), *RAM1* (subunit of the CAAX farnesyltransferase in the protein prenylation branch) and *STE2* (alpha-factor pheromone receptor G-protein). These three genes are directly related to the isoprenoid branch of the mevalonate pathway ^32^. Thus, this small cluster identifies a significant dependence on the isoprenoid enzyme CAAX-farnesyltransferase after deletion of *HMG1, HMG2* and *ARV1* only in the S288C background.

### Dense gene communities are enriched for ER-related functions

Biological networks are generally modular to achieve cellular function and defined by communities ^42,43,44^. Community modules with over-representation of functions define other members of the community also engaged in that function. Using community analysis established for social networks ^45^, we identified 7-14 communities of genes in our 20 GINs (Figure 4). The community panels of the four genetic backgrounds show marked variation in number, density and size as well as the GO term and gene constituents of communities (Supplementary Tables S7-S8). Because the SAFE analysis showed functional enrichment of early secretory pathway processes (Figure 2C), we asked whether there were significant enrichments in GO functions and associated genes in the community analysis. Indeed, in all community panels (Figure 4), there was invariably one community with a characteristic pattern that occurred in dense communities where the GO term *endoplasmic reticulum* (*ER*) displayed high coverage (11-21%, *P*<0.05) and was accompanied by *Golgi apparatus*, *Golgi vesicle transport* and often by *lipid metabolic process*, *protein targeting*, *protein glycosylation* and *cytoplasmic vesicles*.

**Figure 4.**
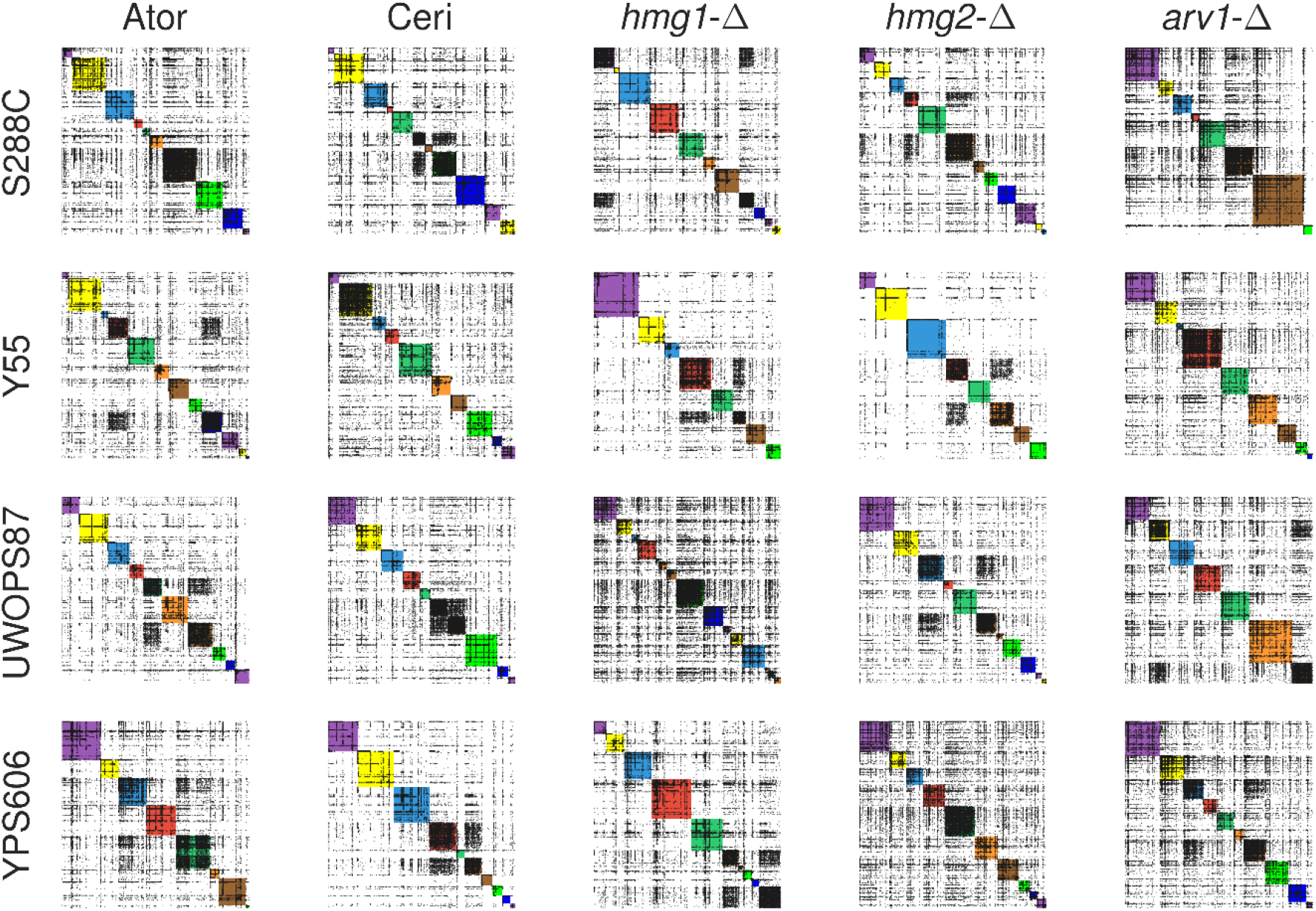
Community analysis reveals genetic background variation among GINs. Communities in each GIN were defined using community analysis previously established for social networks^45^ and coloured on the diagonal and numbered from zero at the top left where each GIN consists of between 7-14 communities (Supplementary Table S8). For example, the GIN in the first panel shows 10 communities.

In more exemplified detail, communities number from top left of each panel starting at #0 and number down the diagonal (Figure 4). Community #6 in the S288C/atorvastatin combination contained this pattern (*ER* function at 12.7%) and is the largest and most dense. Community #4 in the UWOPS87-2421/atorvastatin panel also contained this pattern (*ER* function at 14.5%). Recovery of dense ER function communities with this pattern were also identified in Y55/cerivastatin (community #1, *ER* function at 12.3%), UWOPS87-2421/*hmg1-∆* (community #7, *ER* function at 11.7%), YPS606/cerivastatin (community #5, *ER* function at 11.4%), YPS606/cerivastatin (community #2, *ER* function at 21%), YPS606/atorvastatin (community #2, *ER* function at 19%), Y55/atorvastatin (community #8, *ER* function at 12.8%), S288C/cerivastatin (community #8, *ER* function at 12.7%); S288C/*hmg2-∆* (community #5, *ER* function at 14.6%), S288C/*arv1-∆* (community #0, *ER* function at 11.0%), Y55*/arv1-∆* (community #3, *ER* function at 11.8%), YPS606/*arv1-∆* (community #3, *ER* function at 15.4%), and YPS606/ *hmg2-∆* (community #4, *ER* function at 12.3%). In contrast, another pattern characterised by low *endoplasmic reticulum* term coverage (<5%) was never accompanied by high coverage ER GO term pattern but was almost invariably accompanied by the term *vacuole*, therefore, characterising a different community.

To further investigate the *endoplasmic reticulum* term coverage, we searched for specific genes in each community for membership to this GO term pattern (Supplementary Table S8). *CWH41*, an unfolded protein sensor, was usually found in the most dense community with the high (>10%) *endoplasmic reticulum* coverage regardless of genetic background, suggesting *CWH41* is robustly integral to the UPR. More generally, when GO term communities were interrogated for known early secretory pathway genes such as *CUE1, HOC1, CWH41, CNE1, SIL1, LHS1, KAR2, SEC66, SEC61, IRE1* and *EMC1-6* ^32^, these genes mostly were found in the communities with high *endoplasmic reticulum* coverage. In some cases, other communities of a particular strain/query combination had high *endoplasmic reticulum* term coverage but different ER/Golgi related terms dependent on genetic background. This is an important point because the main recurring pattern establishes a function of the community and though all the other genes in that community are contributing to that function, they can be used differently in different communities, as is apparent by the strain background dependency

### Topology analysis elucidates master regulators of networks

Biological networks have defined topological properties ^47,46 4646^and elucidating network topological metrics adds independent information to functional information about GINs ^14,42^. We calculated three topology measures namely, betweenness centrality (BC; number of shortest path bridging connections to other nodes), closeness centrality (CC; average shortest distance of a node to all other nodes) and eigenvector centrality (EC; a measure of the influence of a node in a network) ^47^. From the distributions of these measures, by genetic background and by query (Supplementary Figures S12-S14), we deconvoluted the genes and identified the five highest betweenness, closeness, eigenvector value genes (Figure 5A; Supplementary Figures S15-S16). These genes by definition are major contributors to topology within our GINs and were strain dependent. For example, *SLT2* highly ranked in the atorvastatin-treated UWOPS87-2421 strain was not highly ranked in other background/query combinations. Similarly, the ER folding sensor *IRE1* had high betweenness scores in the Y55 GINs from atorvastatin and cerivastatin treatment but had markedly (~100-fold) lower scores in UWOPS87-2421 and S288C. The genes that exhibited the high betweenness scores (Figure 5A) are notable because their removal causes the entire network to collapse; these genes are pivotal to the network integrity and as such we deem them *master regulators*.

**Figure 5.**
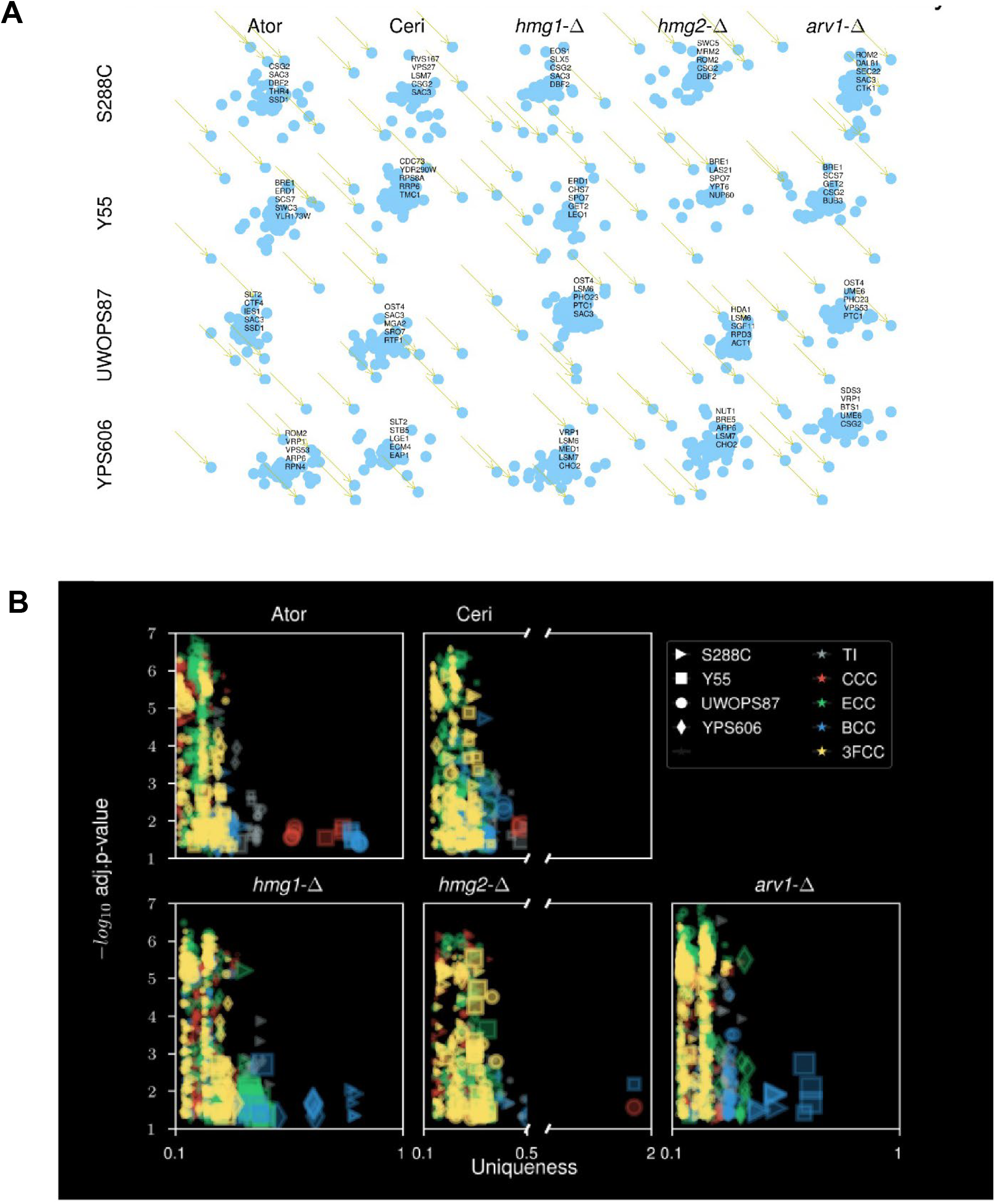
Topology analyses of augmented GINs identify master regulators and functional uniqueness of GINs. A) Deconvoluted master regulator genes with maximum betweenness centrality distinguishes master regulators for each GIN. The top five master regulators are shown for each GIN. B) Gene Ontology enrichment analysis of master regulator genes using multiple topology measurements: TI (topology independent); CCC (closeness centrality cluster); ECC (eigenvector centrality cluster); BCC (betweenness centrality cluster); and 3FCC (closeness, eigenvector and betweenness clusters combined). Shown here are uniqueness (a score that represents the negative of average semantic linearity of a GO term to all other terms), coverage (represented by size of the symbol) and associated statistical significance for each genetic background. For comparison, values were also generated from primary, non-augmented GINs.

### Optimised clustering of network genes confirms functional and community assignments

Although community analysis reveals important network modularity topology features, it may suffer from scalability bias that skews outcomes such as too few large communities and too many small ones caused by artificial fusions or fragmentation of natural clusters, respectively ^48,49^. We therefore tested other conventional clustering heuristics for corroboration. From our augmented GINs we optimized the number of clusters (3-4) for all centrality metrics (3FC, BC, CC, and EC) based on Silhouette clustering score ^49^ and Calinski-Harabasz Index (Supplementary Figure S17; Supplementary Table S9). We named the four clusters α, β, γ and δ which we displayed on a 3-D plot by closeness, eigenvector and betweenness centralities. This plot showed maximum sensitivity to betweenness centrality values (Supplemental Figure S18), demonstrating the fundamental importance of BC to our networks and supporting the similar conclusion reached by the distribution analysis (Figure 5A).

Next we calculated uniqueness values ^50^ from our augmented GINs, after rationalisation of semantically redundant GO terms by a Lin’s similarity pairwise analysis (Supplementary Table S10). We identified a preponderance of betweeness centrality cluster genes in the various query/strain combinations and a lesser number of closeness centrality cluster genes, while the other topology terms were not distinguished (Figure 5B; Supplementary Table S10). There was unique enrichment using betweeness centrality clustering in YPS606/*hmg1-∆* for *mRNA processing* (uniqueness = 0.65), in S288C/*arv1-∆* for ER/*Golgi* (uniqueness = 0.55), and in UWOPS87-2421/atorvastatin for *ribosome processing* (uniqueness = 0.8). These terms, gathered from network topology analysis, show a remarkable correspondence with similar terms in the SAFE result (Figure 2C) that was based only on network function but also corroborate patterns seen in the community analysis.

### Statin-induced UPR is reduced in the statin-resistant strains

The requirement for *IRE1* in statin-resistant strains (Figure 2A-B), led us to directly evaluate UPR activation across the three statin-resistant backgrounds and the statin-sensitive S288C background. UPR measured via a 4X-UPRE element promoter fused to the GFP gene ^37^, did not show significant induction of the UPR in the statin-resistant Y55 and UWOPS87-2421 resistant strains in contrast to the statin-sensitive S288C strain based on confocal microscopy (Figures 6A,6D) and flow cytometry (Supplementary Figure S19). Likewise, the low concentration of cerivastatin induced UPR only in S288C but not in the statin-resistant strains (Figures 6A,6C). In contrast, the high concentration induced UPR in all the strains (Figures 6A,6C), a result consistent with increased side effects of cerivastatin.

**Figure 6.**
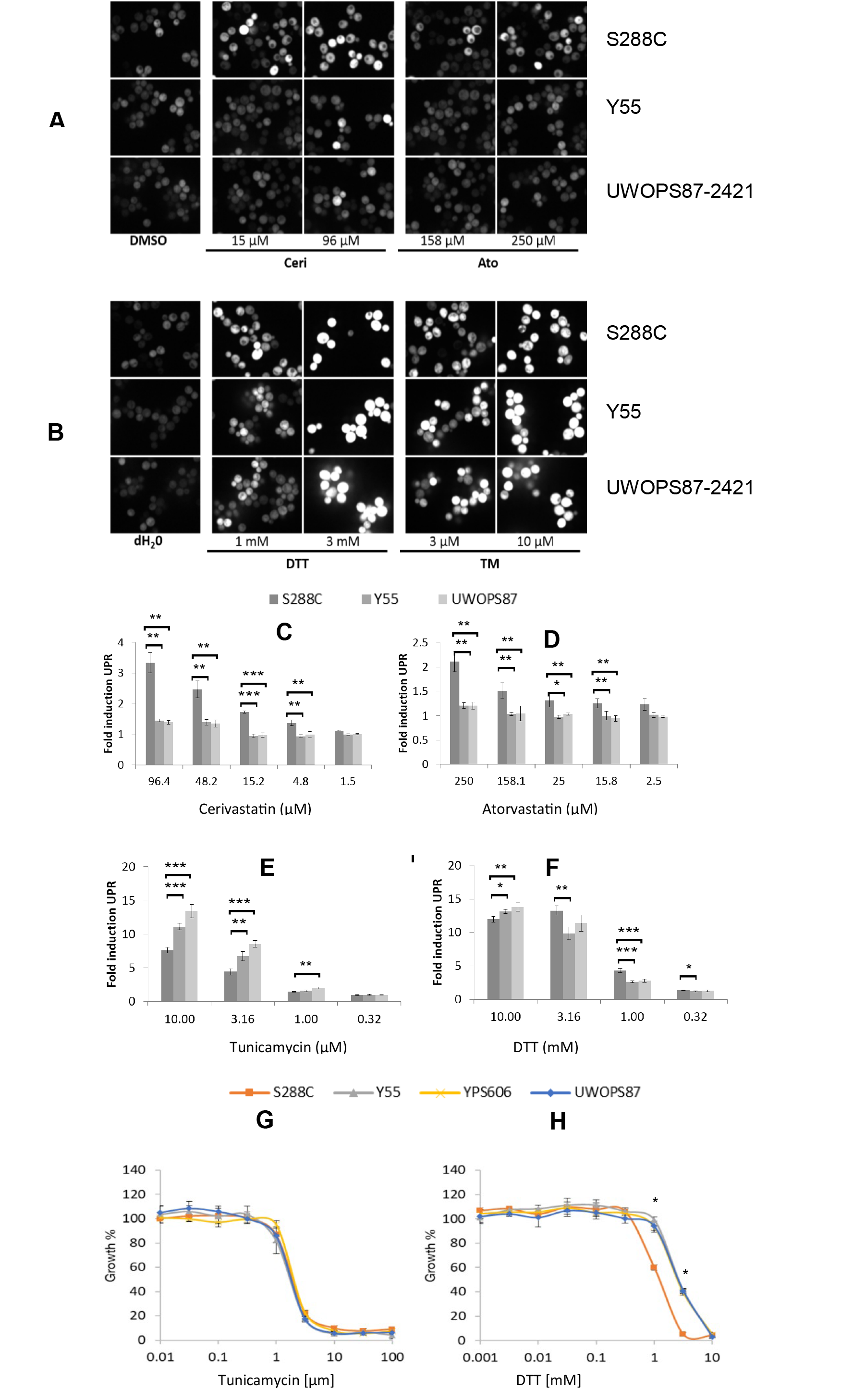
The unfolded protein response mediates statin drug response. A-B) Representative fluorescent microscopy images of mid-log cells for the wild-type strain of S288C, Y55 and UWOPS87-2421 (YPS606 was excluded due to increased flocculation impairing image analysis) expressing 4XUPRE-GFP cells and treated for 4 hours with A) atorvastatin or cerivastatin, or alternatively B) the established UPR inducers dithiothreitol (DTT) or tunicamycin (TM). DMSO and water are the vehicle controls for DTT and TM, respectively. C-F). Fold-induction of UPR in response to C) cerivastatin, D) atorvastatin, E) TM and F) DTT as quantified by the median GFP intensity per cell (n = 200-700 cells) for quadruplicate experiments. *, *P* < 0.05; **, *P* < 0.001; ***, *P* < 0.001 G-H) Growth of the wild-type strain of each genetic background in response to G) TM or H) DTT as shown by percent growth in treated and untreated cells. *, *P* < 0.05, Student’s t-test comparison with S288C.

UPR was readily induced in all the strains with the well-known UPR inducers dithiothreitol (DTT) (Figures 6B,6F) and the N-glycosylation inhibitor tunicamycin (TM) (Figures 6B,6E), showing that the UPR response was not defective in any of the strains. Differential UPR activation in resistant strains was further supported with a growth analysis in which the statin-resistant strains were about a half-log more growth resistant to DTT (Figure 6H) than the statin-sensitive S288C strain but TM equally inhibited growth of all strains (Figure 6G). TM is a PDR substrate ^51^ suggesting the strains are not differentially statin resistant owing to PDR mechanisms. The 4X-UPRE was further quantified using flow cytometry, which corroborated that S288C had increased UPR induction in response to statin treatment relative to the resistant strains (Supplementary Figure S19). Deletion of *IRE1* abolished both tunicamycin and statin induced UPR induction in all strains (Supplementary Figure S19). We conclude that the protein folding system is more robust in the statin-resistant strains than the statin-sensitive S288C strain.

## DISCUSSION

This study aimed to better understand the function of GINs across different strains. Via topology and community analyses of primary and augmented GINs, we defined for the first time genetic background-dependent “master regulator” genes that were required for network integrity. This was demonstrated with the example of atorvastatin and cerivastatin that share a drug target, yet clearly elicit pleiotropic network effects that vary by genetic background among individual yeast strains. Our results highlight the importance of considering GINs in addition to genomics in personalized medicine.

There are a number of innovations in network topological analyses presented in this paper. Analysis of GO terms and genes by communities (Figure 4) and cluster topology (Figure 5) have not been previously conducted for GINs in yeast. We determined that betweenness topology clusters were the most beneficial for network integrity (Figure 5). Impressively, the community analysis is capable of providing nuanced descriptions of different networks arising from the use of different but related genes in closely related cellular processes, but also by using the same genes for different purposes in other communities. This paper identifies network regulators that are only revealed in topology analyses identifying key genes and processes that are not necessarily evident by traditional biochemical analysis of pathways or enrichments of gene lists. An illustration is that the primary functional GINs (Figure 2) featured genes involved with early secretory pathway functions ^37,52^ such as glycosylation, protein folding, ER-Golgi transport, where such patterns were re-capitulated from inspection of the topological community data (Figure 4). A specific illustration is CSG2, an ER protein involved in sphingolipid synthesis, is defined functionally (clusters strongly with *HMG1* Figure 3C) but its high-ranking betweenness score defines it topologically as a network integrity hub in statin response. We also note that this topological property has a genetic background dependency as it appears only in the susceptible strain.

Dependence on genetic background for phenotypic effects has been shown in other studies, though not by comparing GINs. Examples include quantitative trait nucleotides for sporulation efficiency that were dependent on genetic background in 13 yeast strains ^53^, stress QTL that varied across four different yeast strains ^54^, and QTL for ketoconazole and benomyl response that varied across four advanced intercross yeast lines ^55^. Further examples were dependence of epistatic genetic interactions on the genetic background in genome-wide modifier screens ^3^ and another study concluded that network functional modules of protein-protein interactions may be conserved between *Saccharomyces* species, but with substantially different “wiring” ^18,56^. Our study compared strains of different genetic background by GINs derived from SGAs, and a closely similar approach using SGAs to elicit GINs was demonstrated in bacteria in a study on antibiotic resistance ^57^. This study, similarly to ours, concluded that there was very little conservation of GINs, in their case in the response to daptomycin, in two different strains of *Streptococcus pneumoniae* but did not examine topologies, communities and master regulators.

The comparative lack of a UPRE-GFP response to atorvastatin in resistant strains (Figure 6) even though IRE1 is required in the presence of statins (Supplementary Table S4) may be a partial UPRE response ^58^ or a more robust protein folding system in the resistant strains as seen by greater resistance to DTT (Figure 6). We note that the statin susceptible S288C strain shows a UPR response (Figure 6) in the presence of statins, therefore the UPR sensing gene, *IRE1*, must be used, but it is not essential to S288C ^36,52^. Additionally, the ‘hub gene’ *SPF1* that clustered closely with functionally related *IRE1* (Figure 3B), must be causing UPR activation through altered ER lipids, and not directly by the *IRE1* unfolded protein sensing. An explanation is that in atorvastatin treated S288C, ER-stress is mainly buffered by multivesicular body (autophagy) associated genes as seen in the SAFE analysis (Figure 2C) similar to observations reported for *C. elegans* ^59^ and human cells ^60^ whereas in the resistant strains atorvastatin toxicity is buffered by a more robust folding/ERAD system (Figure 6). These observations taken with others in this paper lead to the conclusion that a stress phenotype (ER stress) may be buffered by quite different processes dependent on the genetic background of the strain.

If specific GINs are not conserved across genetic backgrounds, then what are the detailed functions of GINs? GINs have been described as having intrinsic functionality ^61,62^ or as functional modules related to topological modules ^46,63^, and thus one might ask if GINs involving hundreds of genes are units of heritability? Human height involves hundreds of genes of small additive effect genes ^64^ that must be heritable as units because height is a heritable trait. However, we have concluded that GINs cannot be units of heritability because GINs and their involved processes, ER-stress response to statins in this paper, were not conserved in closely related strains. Although GINs appear to be ephemeral or transitory, GINs could nonetheless have a role in evolutionary potential ^57,65^. The concept of an evolutionary ratchet of sequentially accumulating single mutations stabilized by within-protein epistatic effects describing the evolution of the glucocorticoid receptor ^66^ might be instructive. It is thus plausible that GINs emergent on a new mutation are an evolutionary ratchet that irreversibly stabilise a new mutation one at a time allowing increased fitness through transient epistatic mechanisms superseded over time by additional mutations. However, the GINs in this model would not be heritable but their effects would be heritable. Such a model could be tested by observing whether an extant GIN seen in a particular strain deletion library was ephemeral after additional meiosis but stable after additional mitosis.

In summary, we show the statin-resistant strains exhibited a dependence on protein folding and glycosylation genes that was sufficient to alleviate the requirement for the UPR. By contrast, in the statin-susceptible S288C strain, ER stress was evident by induction of the UPR, perhaps buffered by requirement for protein degradation processes through multivesicular bodies, autophagy and the proteosome. This is a nuanced yet critical difference that would not be evident without analysis of GINs in the individual strains. We conclude that GIN function and the topological properties of the network induced by a drug, in addition to the main target, must be considered in the drug mechanism of action and efficacy. The use of deletion libraries in multiple genetic backgrounds will guide future eukaryotic chemical genomic analyses from yeast to human cells.

## METHODS

### Yeast strains and media

*Saccharomyces cerevisiae* strains (Supplementary Table S1) including SGRP strains (National Collection of Yeast Cultures (NCYC), Norwich, United Kingdom) and the S288C DMA library were maintained in synthetic complete (SC), synthetic dropout (SD), enriched sporulation, or yeast peptone dextrose (YPD) as previously described ^67^. Media included yeast extract, peptone, yeast nitrogen base without amino acids and ammonium sulphate, amino acids and agar (Formedium Ltd.), nourseothricin (Werner BioAgents), Geneticin (G418) and Hygromycin B (HPH) (Life Technologies Ltd.), 5-Fluoroorotic Acid (Kaixuan Chemical Co. Ltd.), atorvastatin (Inter Chemical Ltd.), cerivastatin (Chengdu Caikun Biological Products Co. Ltd.) and ampicillin, canavanine and S-aminoethyl-L-cysteine (Sigma).

### Backcross construction of SGRP deletion mutant array (ssDMA) libraries

Backcross methodology was used construct ssDMA libraries. Appropriate selection markers were introduced in SGRP strains via PCR-mediated disruption with a variety of primers (Supplementary Table S11). The KanR marker at the *URA3* locus in SGRP strains was replaced with pAG60-derived *CaURA3 (*Supplementary Table S12) ^68^, which was then removed by homologous recombination with the flanking *ura3Δ0* region from BY4742, followed by growth on 5-FOA media selecting for *URA3* null mutation. The selected SGRP strains were crossed with the original S288C DMA and the hybrid progeny backcrossed five more times with the SGRP strains, resulting in three new deletion libraries we termed “ssDMAs” for SGRP strain deletion mutant arrays.

### Genome sequence analysis and genomic authenticity of ssDMA libraries

The ssDMA strains were subjected to whole genome sequence analysis to determine the efficacy of the backcross strategy. Genomic DNA was extracted from pooled yeast strains from plate 10 of the DMA and ssDMA libraries. Truseq Nano 350 bp insert libraries were prepared (using a TruSeq Rapid SBS kit or TruSeq SBS kit v4 and sequenced on an Illumina HiSeq2000 instrument (Macrogen, South Korea). Raw image processing, base calling, and conversion to FASTQ format was carried out at Macrogen by HiSeq Control Software v2.2 Real Time Analysis v1.18.61 and bcl2fastq v1.8.4. All sequence data were aligned to the *Saccharomyces cerevisiae* S288C reference genome ^32,69^ and SGRP strains^70^. The data alignment pipeline was carried out as previously described ^55^. Whole Genome Vista ^71^ was used to align the FASTA sequence of each ssDMA to the respective parental SGRP strain. The pairwise alignment score was calculated from the summation of the alignment score of each aligned fragment with a weighting on the basis of fragment size and filtration to only retain regions that had 100% alignment between the ssDMA sequence and parental strains. Sequence coverage in the final BAM file was used to determine if any aneuploidy was evident in any of the ssDMA libraries. Per-base sequence coverage was exported using Unipro UGENE v1.27 and the mean average coverage was calculated for each 20 kb genomic bin.

### Synthetic genetic array analysis

SGA analysis was conducted as previously described ^28^ with the additional methodology of integrating SGA reporters in the MATα ssDMA strains. The *CEN-URA3* marker (pRS316) was expressed in the standard SGA query strain (Y7092) with the reporter (*can1Δ::STE2pr-Sp_his5*), mated with the SGRP parent, and the CEN-URA3 marker was removed with selective killing on media containing 5-FOA. Replica plating (pinning) was performed using an automated RoToR had system (Singer Instruments). Double deletion mutants at completion of the SGA analysis were incubated at 30°C for 48 h and imaged using a digital camera (Canon EOS 600D).

### Chemical genetic analysis

Growth of ssDMA and S288C libraries was evaluated concentrations of atorvastatin and cerivastatin which inhibited growth by approximately 40% in SC media. These concentrations of atorvastatin (25, 100 μM) and cerivastatin (10, 50 μM) were based on growth of plate 10 in each library in the screening format of 1536 colonies per plate. The S288C and ssDMA libraries were pinned in 1536 format on the media containing either atorvastatin, cerivastatin or DMSO (vehicle control), incubated at 30°C for 48 h, and imaged using a digital camera (Canon EOS 600D).

### Phenotypic analysis

Colony size and circularity were measured using Gitter^72^ and statistically compared using ScreenMill ^73^. Growth ratios (double deletion *vs* single deletion, treated single deletion *vs* untreated single deletion) were represented in *z*-score values that were statistically evaluated using a normal distribution. Interactions (*Z* > 2.0, *P* < 0.05) were visualized using Cytoscape v3.5.1 ^74^.

### SAFE analysis

SAFE analysis was performed as previously described ^12^ using the global genetic interaction network of *Saccharomyces cerevisiae* ^13,32^, augmented where the map-weighted shortest path length was the distance metric, maximum distance from a node was set at 7.5 radius, and threshold for enrichment significance was 0.05.

### Topology analysis

Closeness centrality, betweenness centrality, and eigenvector centrality were calculated as previously described ^63^ on a global genetic interaction network ^13,75^ that included all genes less than two levels of distance apart (*i.e.*, path length of 3) from the seed vertices (list of gene from the 20 primary SGAs). Interactions were additionally filtered based on established stringent cut offs for positive digenic interactions, digenic negative interactions, and trigenic interactions ^13^. Bootstrap analysis (n = 1000) of a random sample set of genes of the same sizes was statistically compared with our list using a Mann-Whitney rank test. Only networks with *P*-values below 0.05 were included in the topology analyses.

### Functional community clustering analysis

Networks were visualized as sorted adjacency matrices for best modularity using the Louvain method ^76^. Defining *A* as edge weight, *k* as the sum of the weights of the edges attached to respective vertex, *2m* as the sum of all of the edge weights in the graph, *C* is the community, and *δ* as Kronecker delta for a weighted network with vertices *i* and *j*, then random walk modularity *M* is defined as:

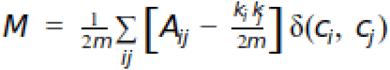

where *A* represent edge weight, *k* is the sum of the weights of the edges attached to respective vertex, *2m* is the sum of all of the edge weights in the graph, *C* is the community, and *δ* is a Kronecker delta. Change of modularity after reassigning the community is calculated as follows:

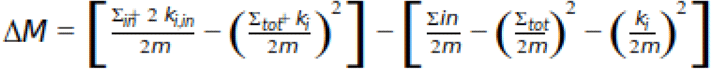

### GO enrichment analysis

Gene Ontology (GO) terms were identified using GOATOOLS ^77^ for each community and statistically evaluated using *P*-values that were corrected for by FDR 2-stage Benjamini-Krieger-Yekutieli. Distinct, functional communities were defined by ‘Uniqueness’, a score that represents the negative of average semantic similarity of a term to all other terms according to Lin’s semantic similarity ^78^.

### Unfolded protein response analysis

The UPR was evaluated using a highly specific fluorescent marker as previously described ^79^. Strains were transformed to express 4XUPRE-GFP comprising four tandem repeats of the UPR elements fused to a GFP. Yeast strains subcultured to an OD of 0.15 were grown for 2 h and treated with appropriate concentrations of UPR-inducing agents (DTT, TM) or statins (atorvastatin, cerivastatin) at 30°C for 4 h. Fluorescent signal was detected at 488 nm (GFP) using a 60x water immersion lens (NA 1.2) in an automated, high-throughput spinning disk confocal microscope (Evo Tec OPERA, Perkin Elmer). Three visual fields at the same exposure were quantified in quadruplicate experiments for each condition using Acapella automated image analysis software (Perkin Elmer) as previously described ^36^.

### Flow Cytometry

4XUPRE-GFP was measured as previously described with modifications ^80^. Cells containing the 4XUPRE-GFP construct were cultured at an OD of 0.2 and grown in atorvastatin, cerivastatin, tunicamycin, or DMSO for 4 h. Cells were stained with 4 μg/mL of propidium iodide immediately prior to processing to determine cell viability. Fluorescence was measured using a FACSanto^TM^ II flow cytometer (Becton Dickinson). Dead cells were excluded from fluorescent measurements. Geometric mean fluorescence and Median Absolute Deviation (MAD) was calculated using FlowJo.

### Data availability

The datasets for this article are available as part of the Supplementary Information. Raw genome sequence data from the three new ssDMAs (Y55, UWOPS87-2421, YPS606) are publically available at SRA number SRP124399 and BioProject PRJNA417352.

## ACKNOWLEDGEMENTS

We acknowledge Professor Charles Boone (University of Toronto) for the kind gift of the S288C deletion collection.

## AUTHOR CONTRIBUTIONS

B.P.B. and P.H.A. conceived the project. B.P.B. created the deletion collections and carried out the experimental work relating to GINs. E.N. carried out the computational SAFE analysis, hierarchical clustering, topology analysis, and community analysis. C.A.R. completed the genome sequence analysis. N.V.C., J.S. and D.S. carried out the experimental work relating to the UPR. P.H.A, B.P.B and A.B.M. wrote the manuscript. P.H.A. supervised the project. All authors reviewed and approved the final version of the manuscript.

---

**Supplementary Figure S1.**
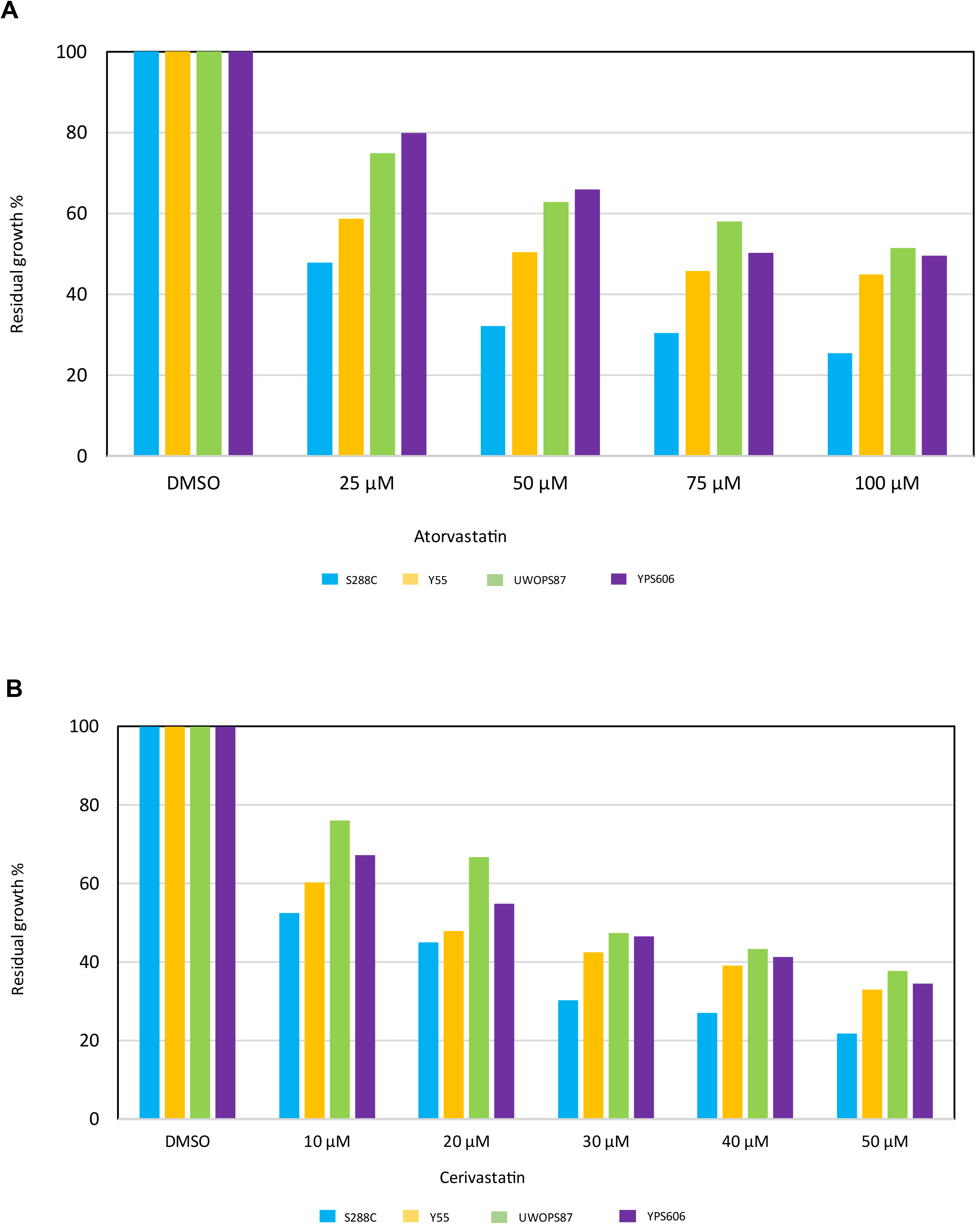
Residual growth (a ratio of growth in treated and untreated cells) of all 1536 colony size values from plate 10 of the original S288C DMA compared to the three newly created ssDMA libraries in the genetic backgrounds of Y55, UWOPS87-2421 (UWOPS87) and YPS606. Growth was quantified in A) atorvastatin and B) cerivastatin.

**Supplementary Figure S2.**
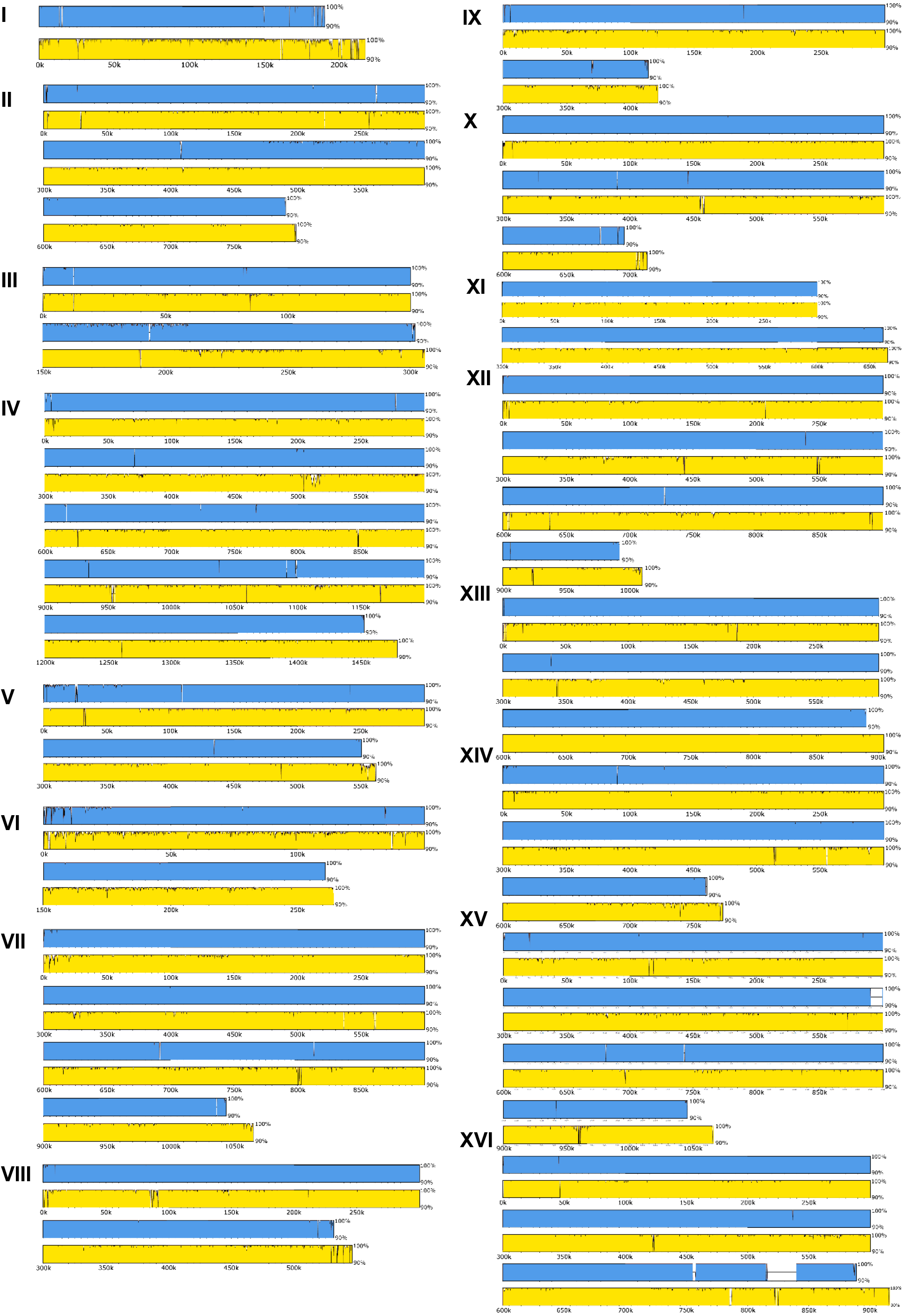
UWOPS87-2421 ssDMA alignment to UWOPS87-2421 parental strain (blue) and S288C parental strain (yellow).

**Supplementary Figure S3.**
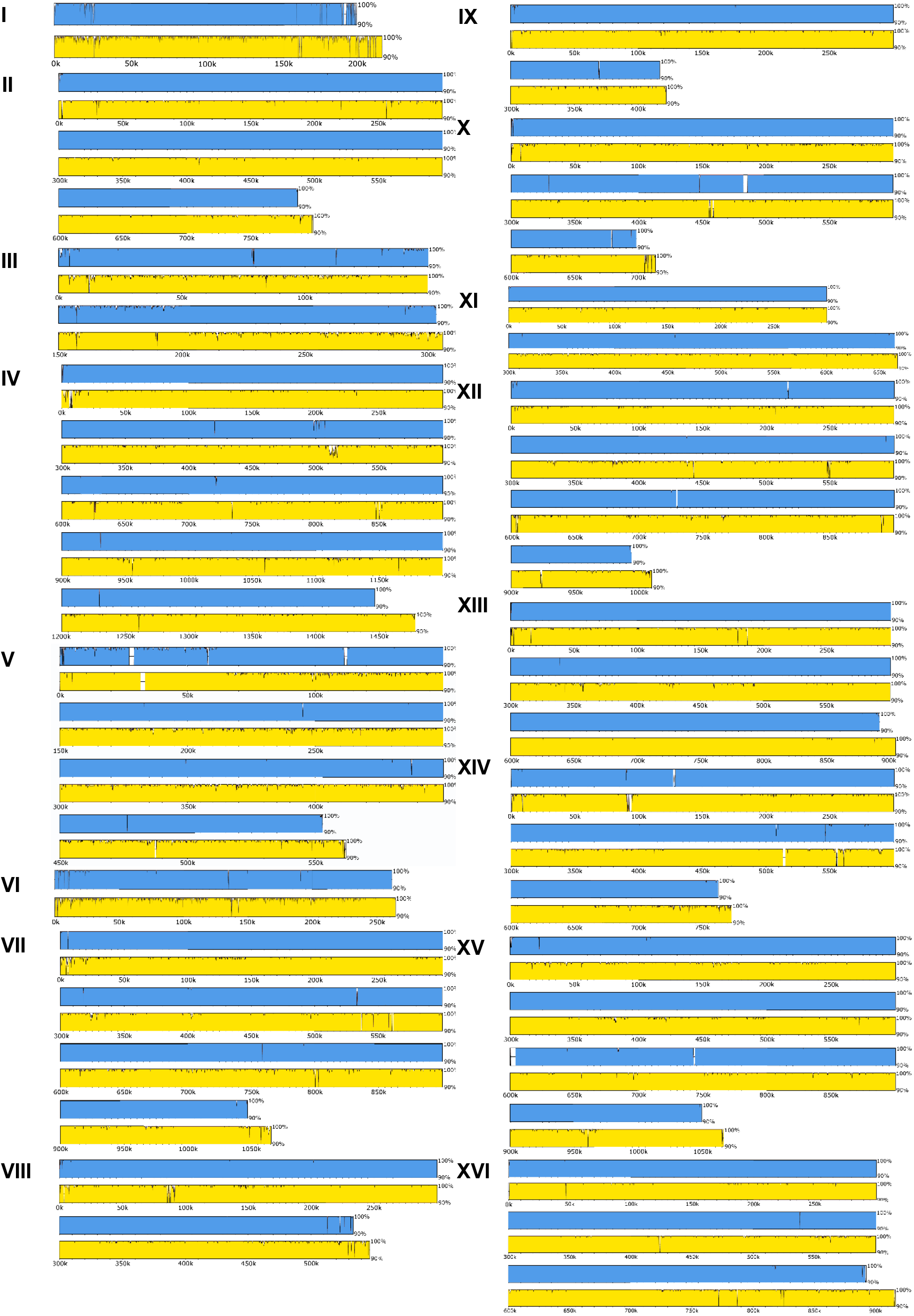
Y55 ssDMA alignment to Y55 parental strain (blue) and S288C parental strain (yellow).

**Supplementary Figure S4.**
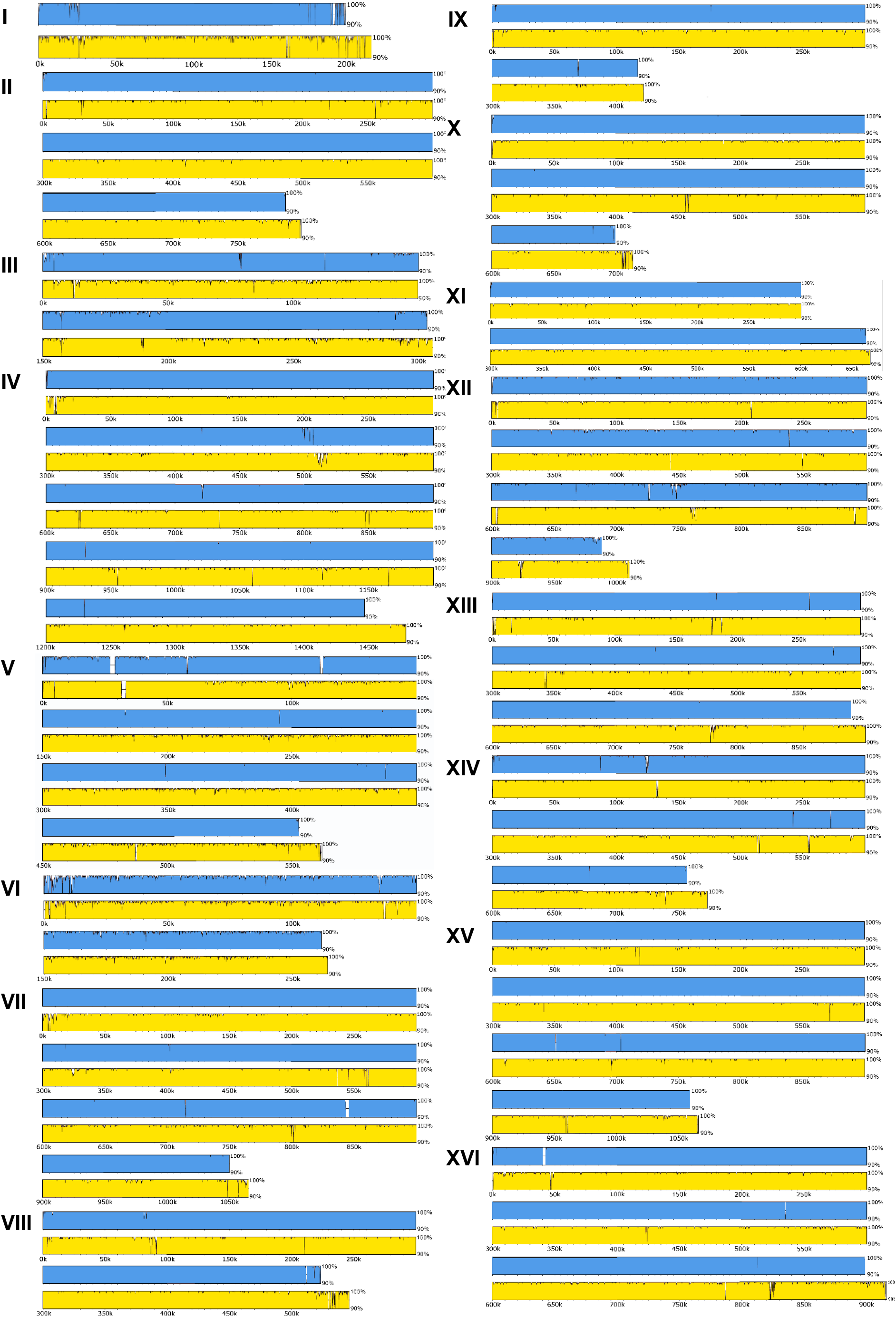
YPS606 ssDMA alignment to YPS606 parental strain (blue) and S288C parental strain (yellow).

**Supplementary Figure S5.**
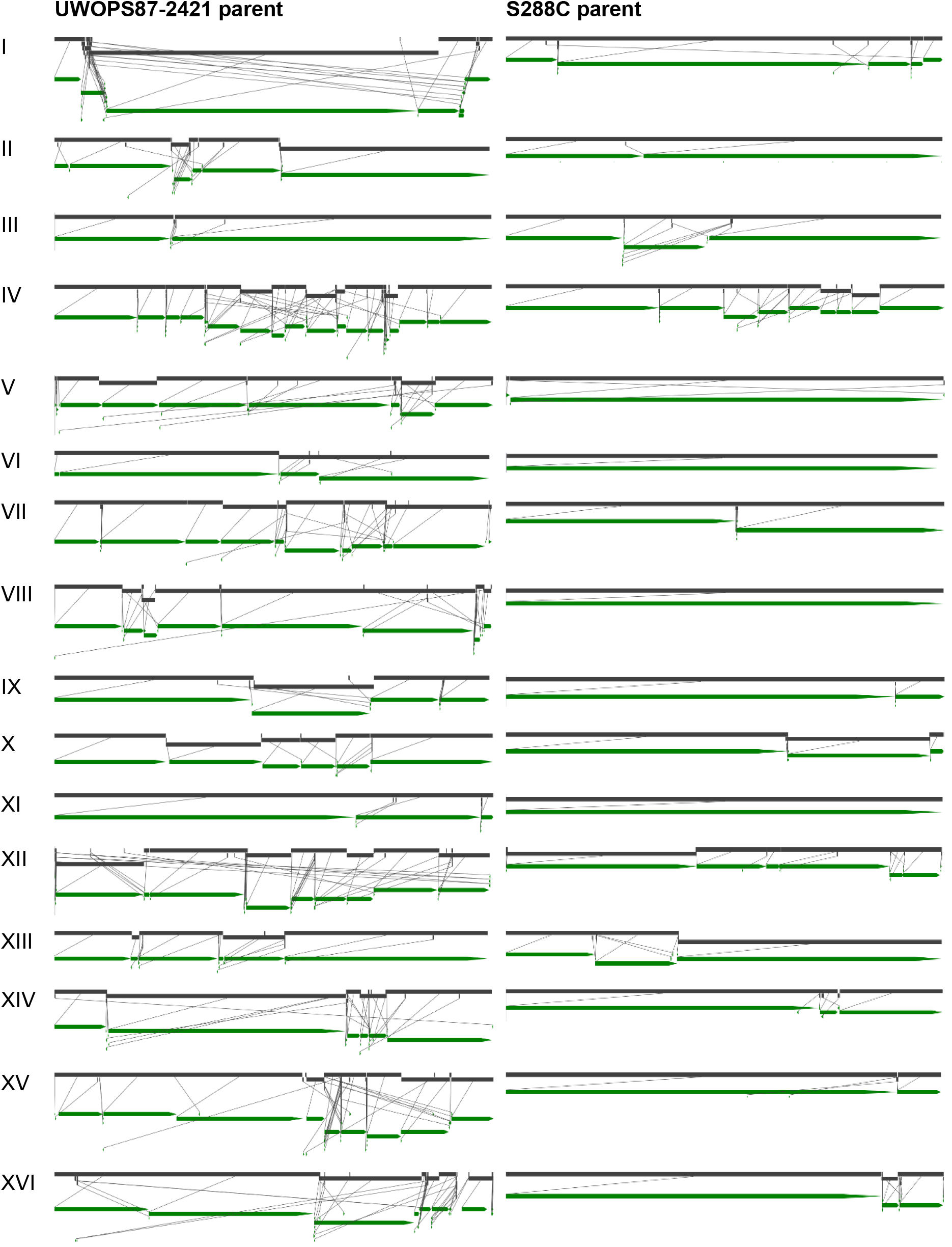
Synteny between UWOPS87-2421 ssDMA (grey) and its respective UWOPS87-2421 and S288C parental strains (green).

**Supplementary Figure S6.**
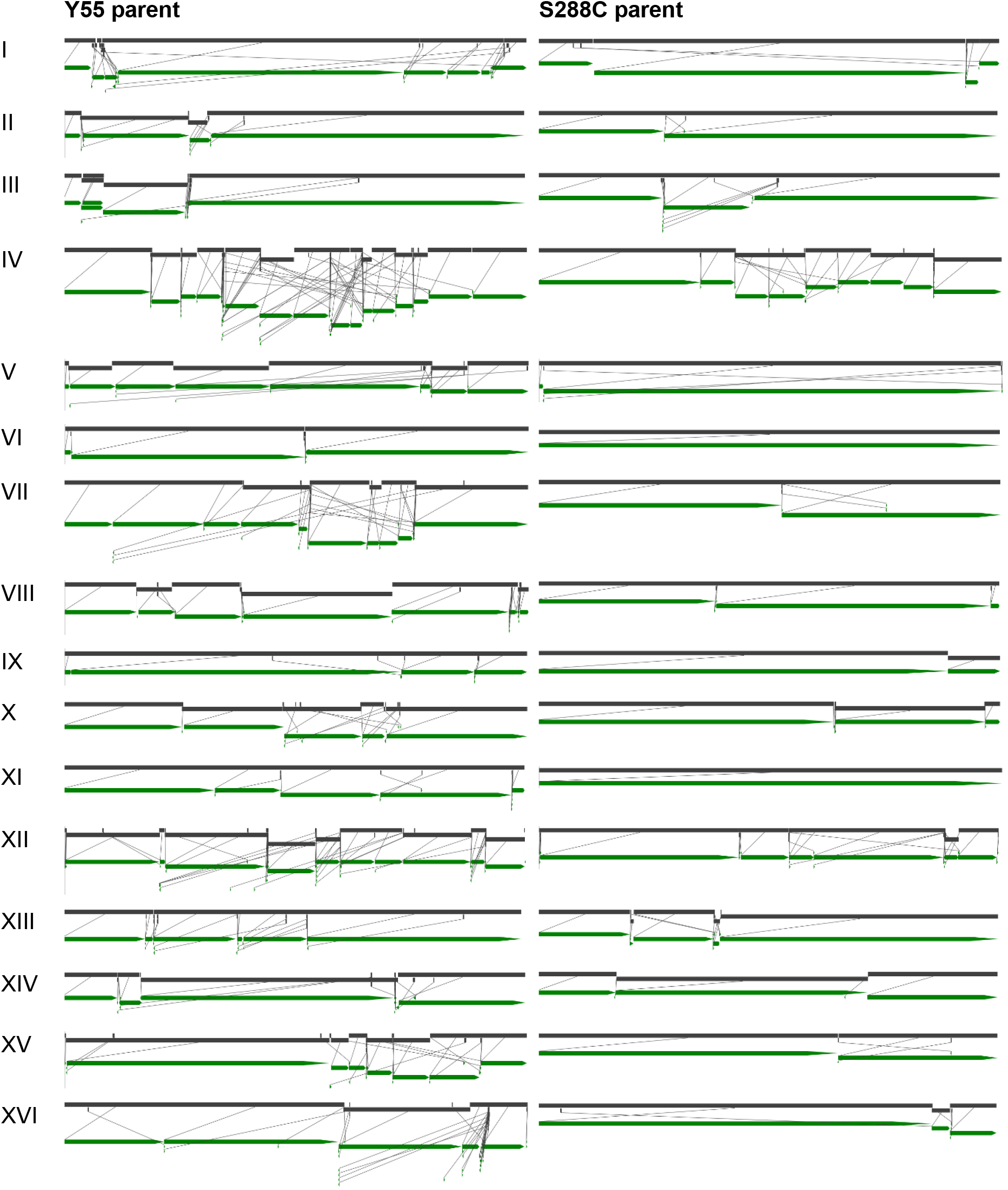
Synteny between Y55 ssDMA (grey) and its respective Y55 and S288C parental strains (green).

**Supplementary Figure S7.**
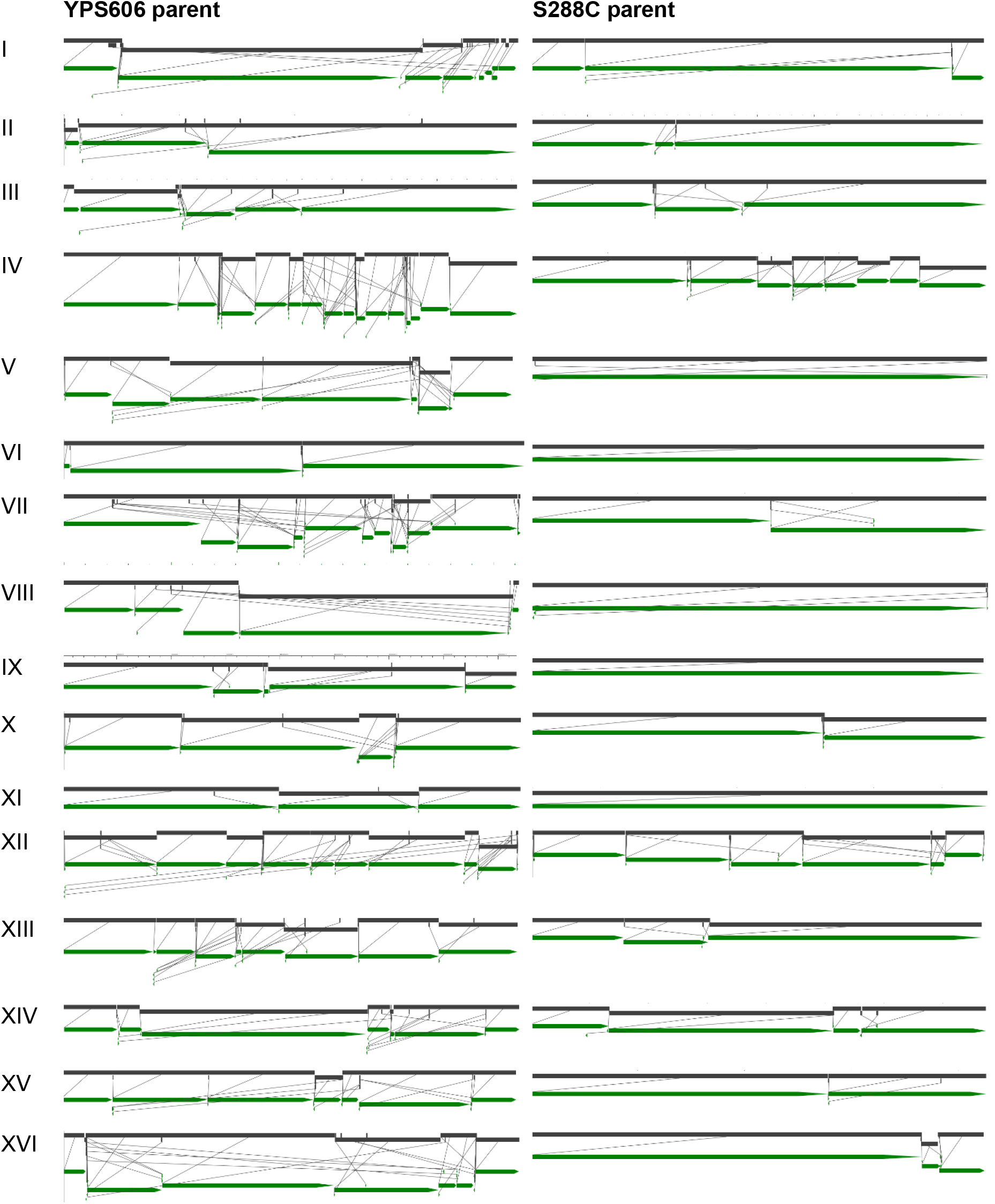
Synteny between YPS606 ssDMA (grey) and its respective S288C and S288C parental strains (green).

**Supplementary Figure S8.**
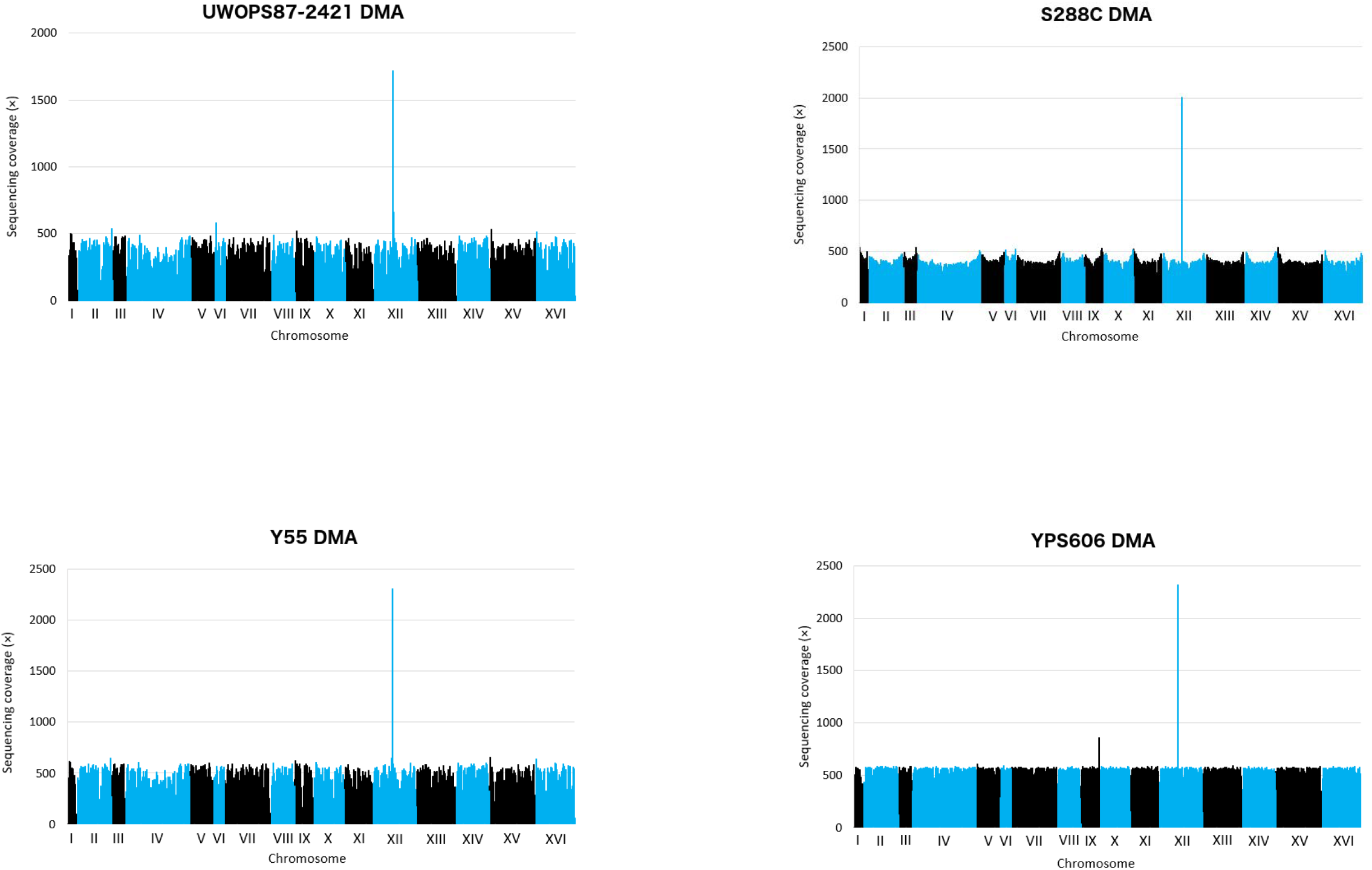
Sequencing coverage of the ssDMA libraries (UWOPS87-2421, Y55, YPS606) and the control S288C DMA library shows no evidence of chromosomal aneuploidy.

**Supplementary Figure S9.**
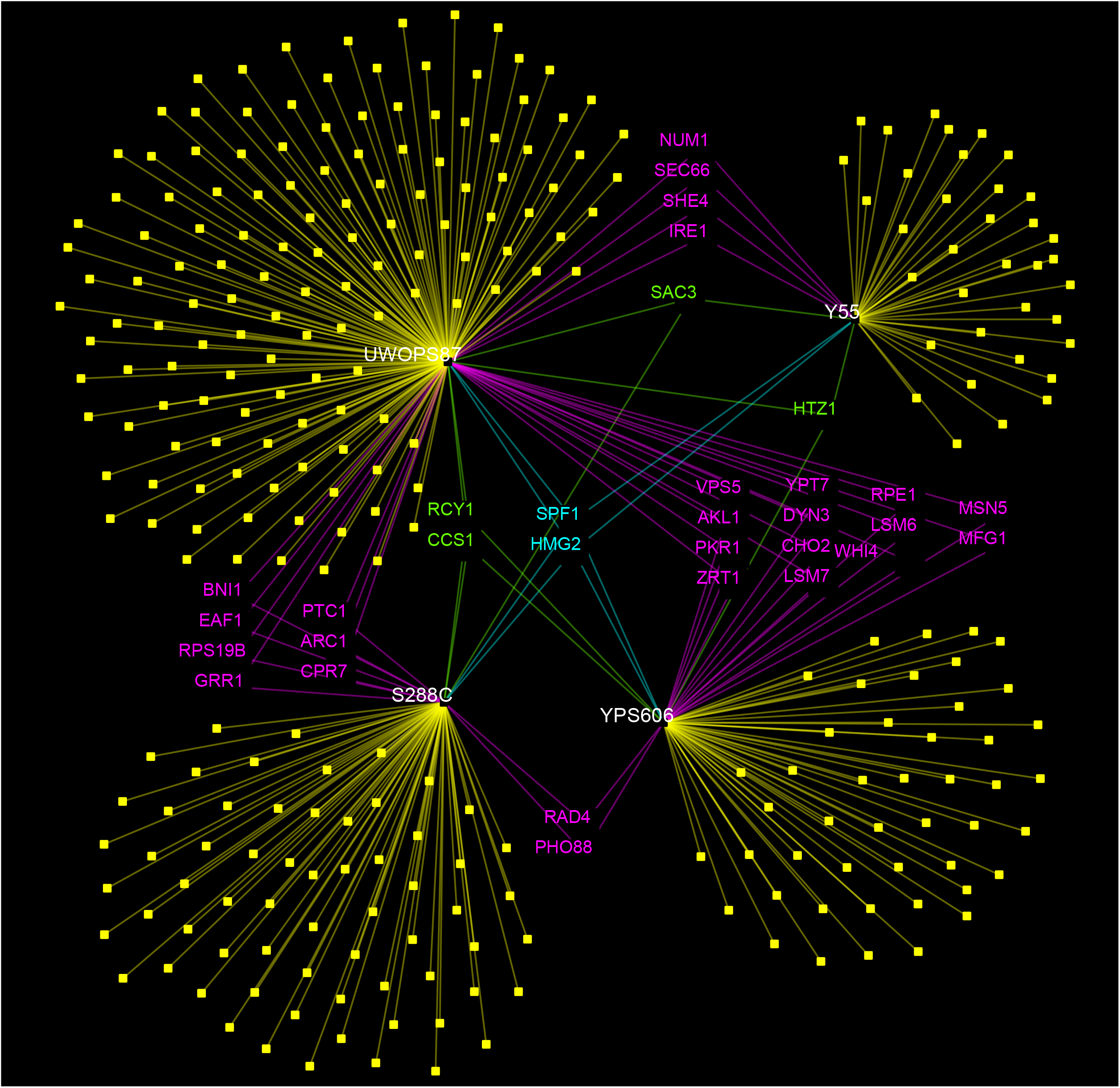
Primary genetic interaction network representing *HMG1* interactions in the four genetic background deletion libraries (S288C, Y55, UWOPS87-2421 (UWOPS87) and YPS606). Interactions are shared by different genetic backgrounds (purple, green, and blue) or unique to specific genetic backgrounds (yellow, genes not included in this figure).

**Supplementary Figure S10.**
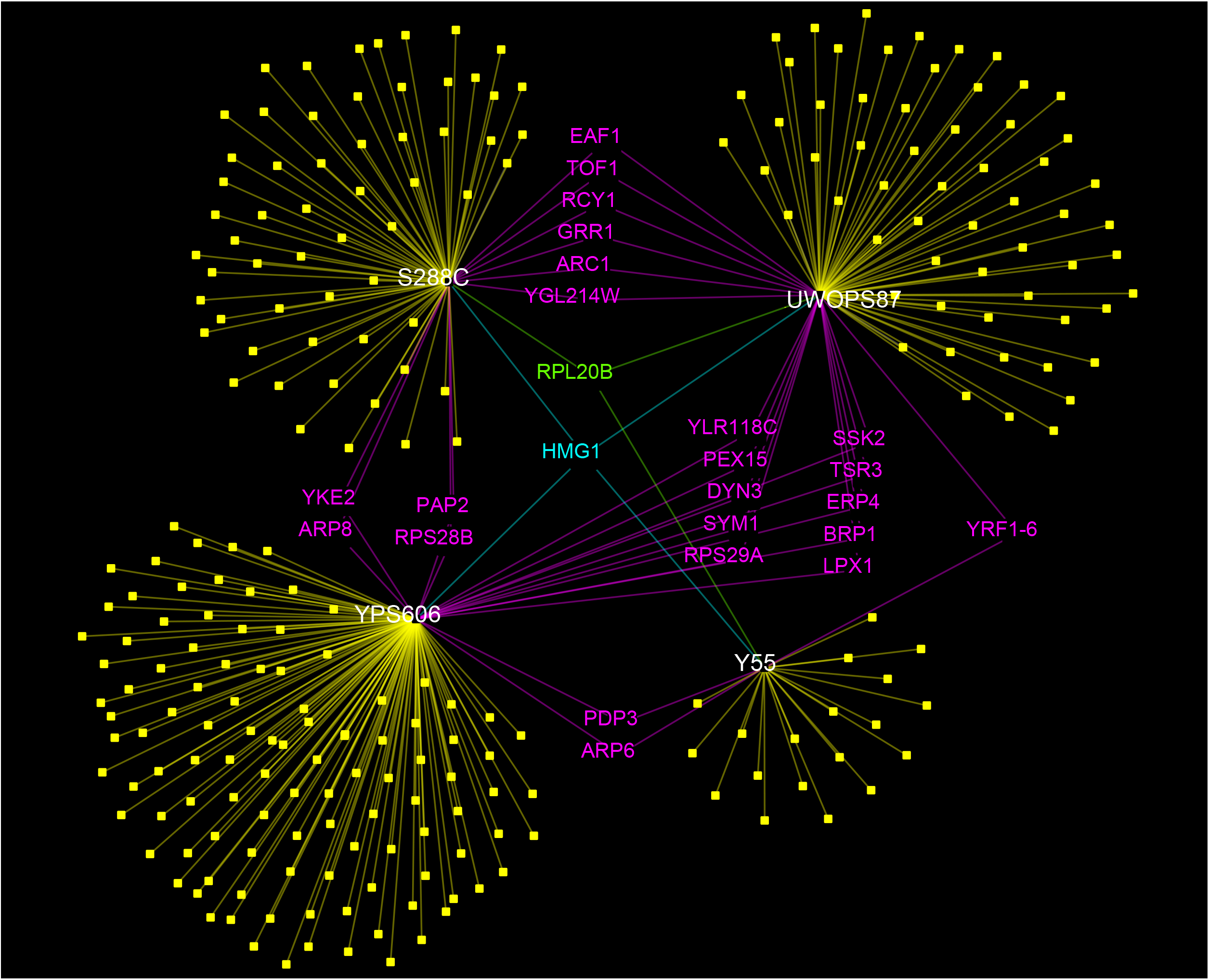
Primary genetic interaction network representing *HMG2* interactions in the four genetic background deletion libraries (S288C, Y55, UWOPS87-2421 (UWOPS87) and YPS606). Interactions are shared by different genetic backgrounds (purple, green, and blue) or unique to specific genetic backgrounds (yellow, genes not included in this figure).

**Supplementary Figure S11.**
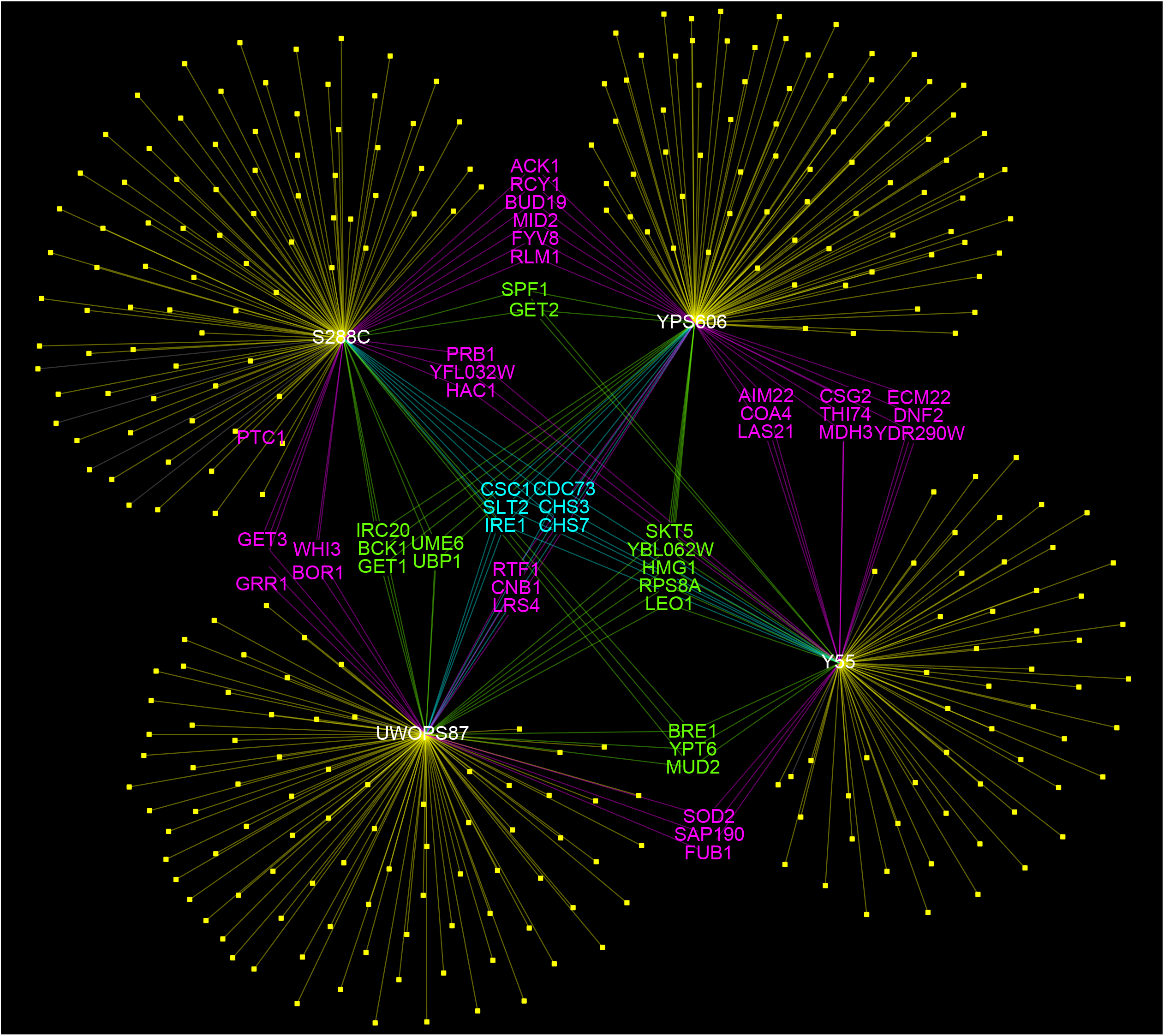
Primary genetic interaction network representing *ARV1* interactions in the four genetic background deletion libraries (S288C, Y55, UWOPS87-2421 (UWOPS87) and YPS606). Interactions are shared by different genetic backgrounds (purple, green, and blue) or unique to specific genetic backgrounds (yellow, genes not included in this figure).

**Supplementary Figure S12.**
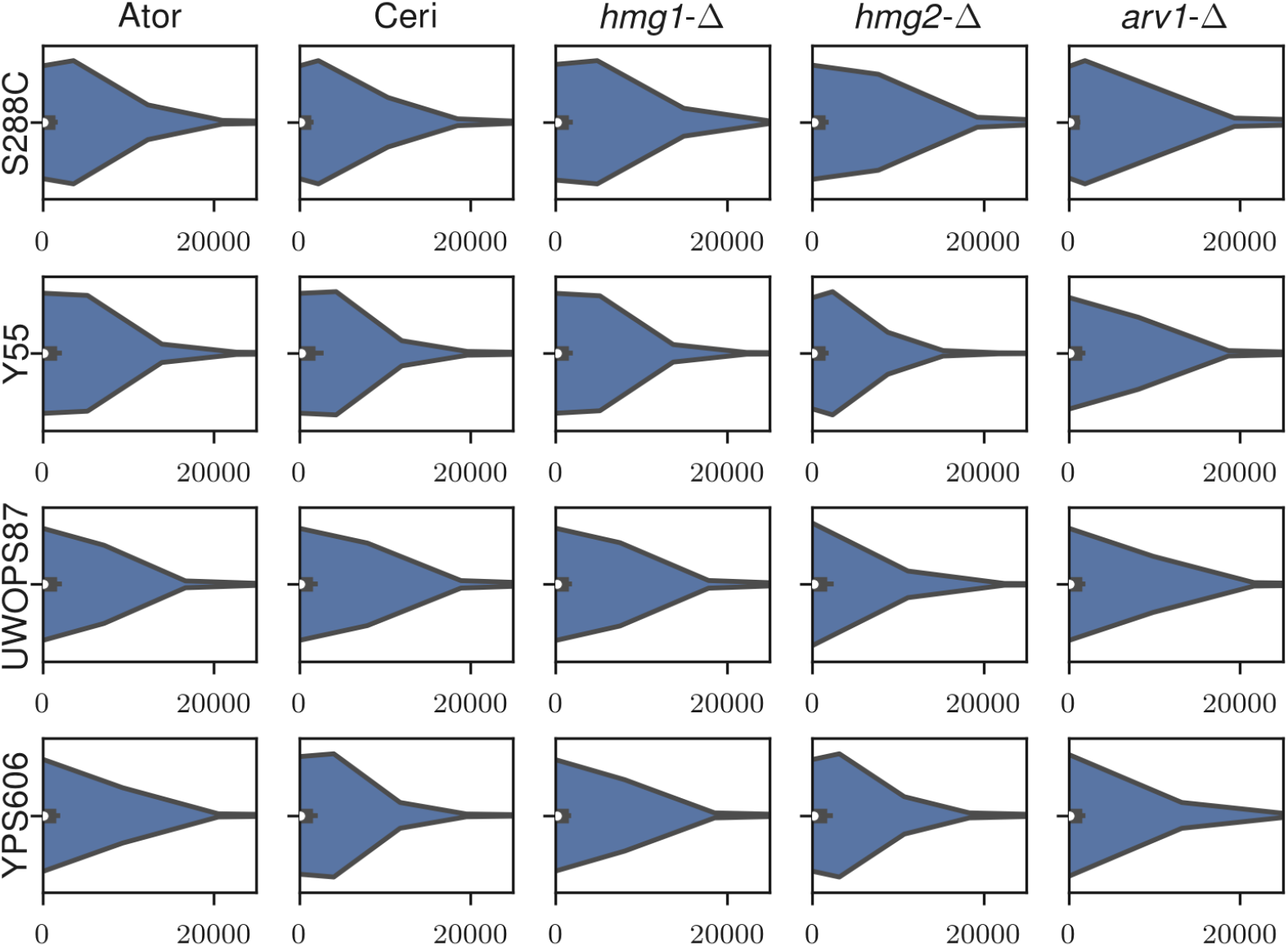
Distribution of betweenness centrality for DMA libraries in four strains (S288C, Y55, UWOPS87-2421 (UWOPS87) and YPS606) by five queries (atorvastatin, cerivastatin, *hmg1-Δ, hmg2-Δ* and *arv1-Δ*).

**Supplementary Figure S13.**
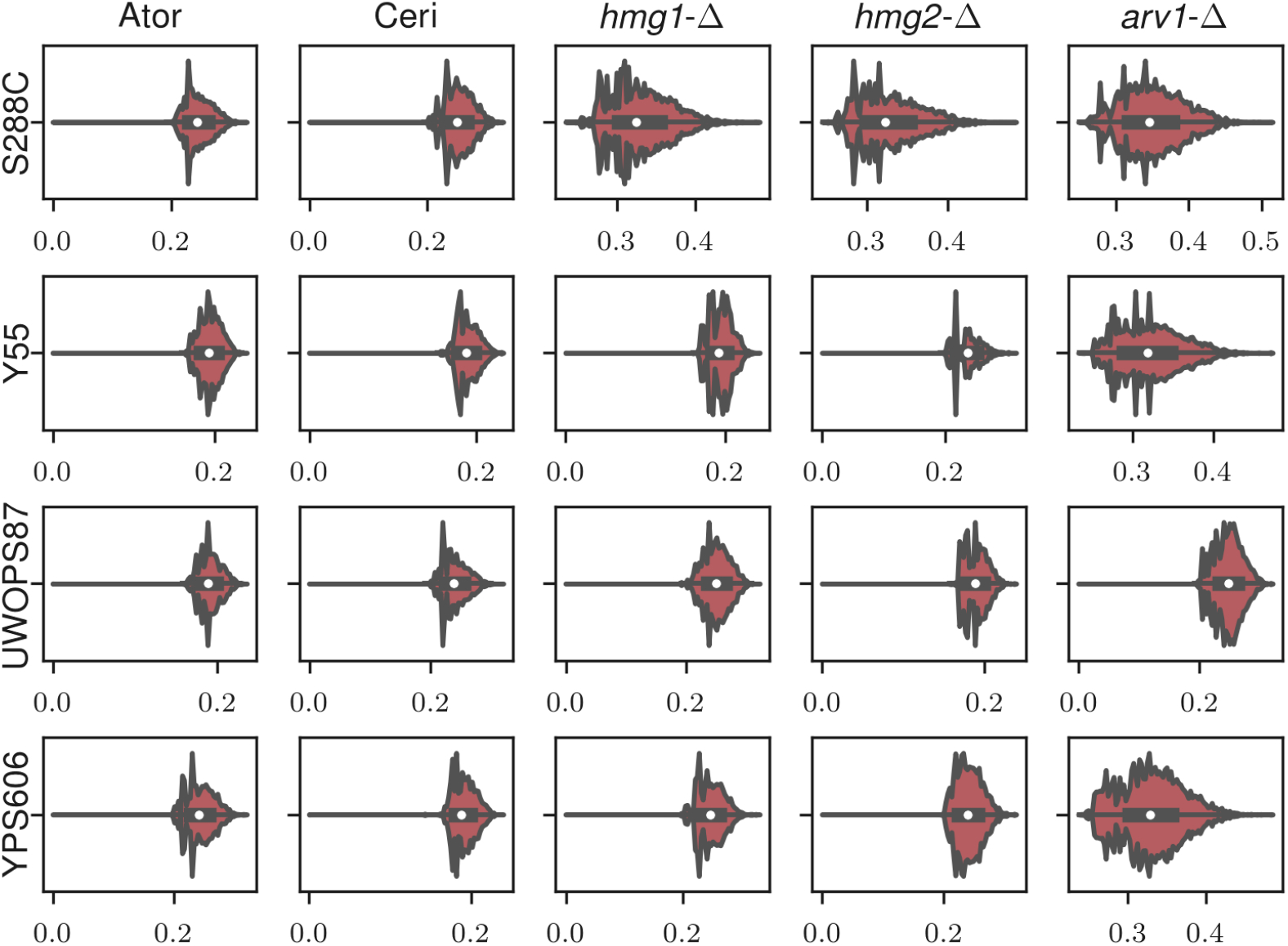
Distribution of closeness centrality for DMA libraries in four strains (S288C, Y55, UWOPS87-2421 (UWOPS87) and YPS606) by five queries (atorvastatin, cerivastatin, *hmg1-Δ, hmg2-Δ* and *arv1-Δ*).

**Supplementary Figure S14.**
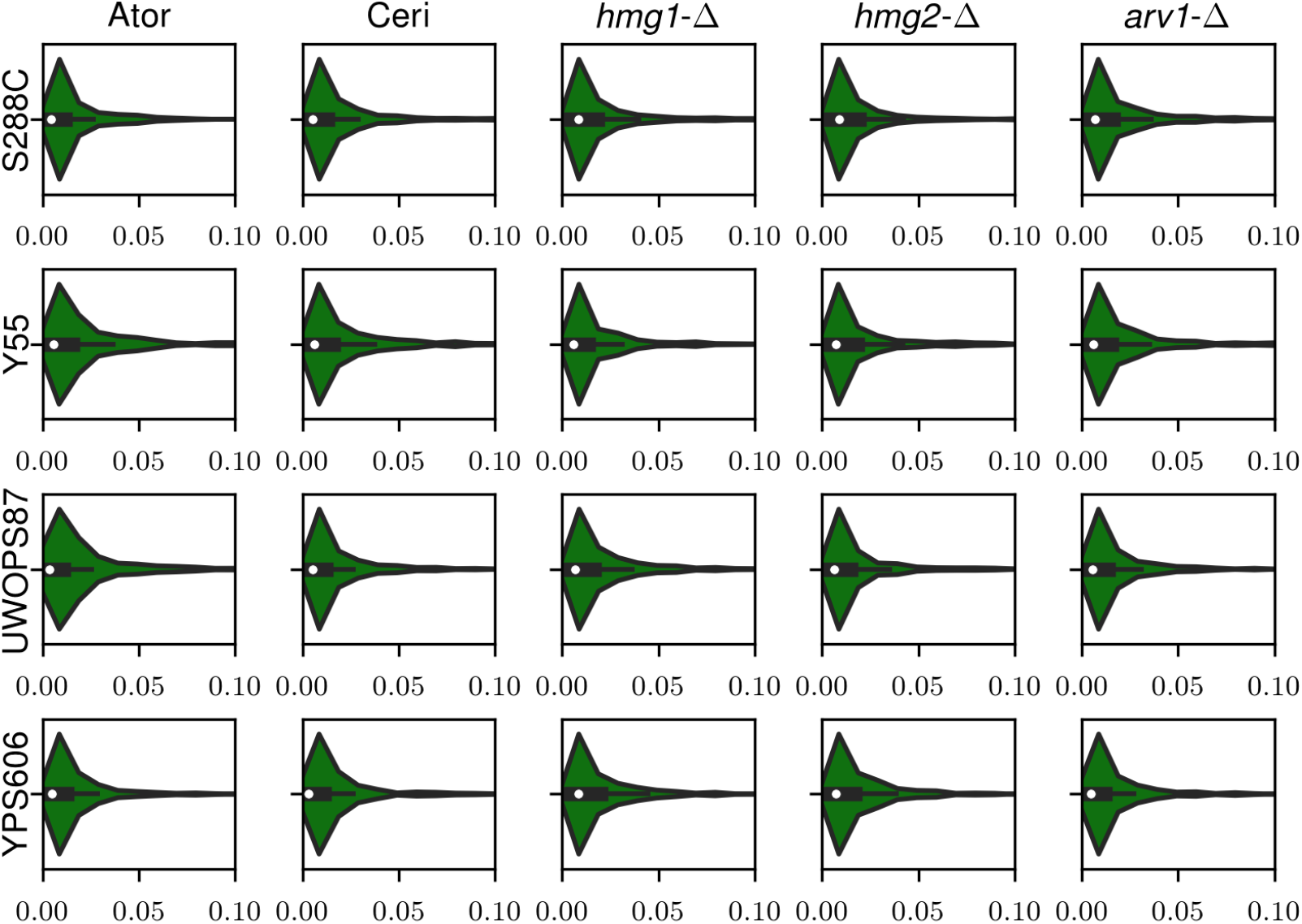
Distribution of eigenvector centrality for DMA libraries in four strains (S288C, Y55, UWOPS87-2421 (UWOPS87) and YPS606) by five queries (atorvastatin, cerivastatin, *hmg1-Δ, hmg2-Δ* and *arv1-Δ*).

**Supplementary Figure S15.**
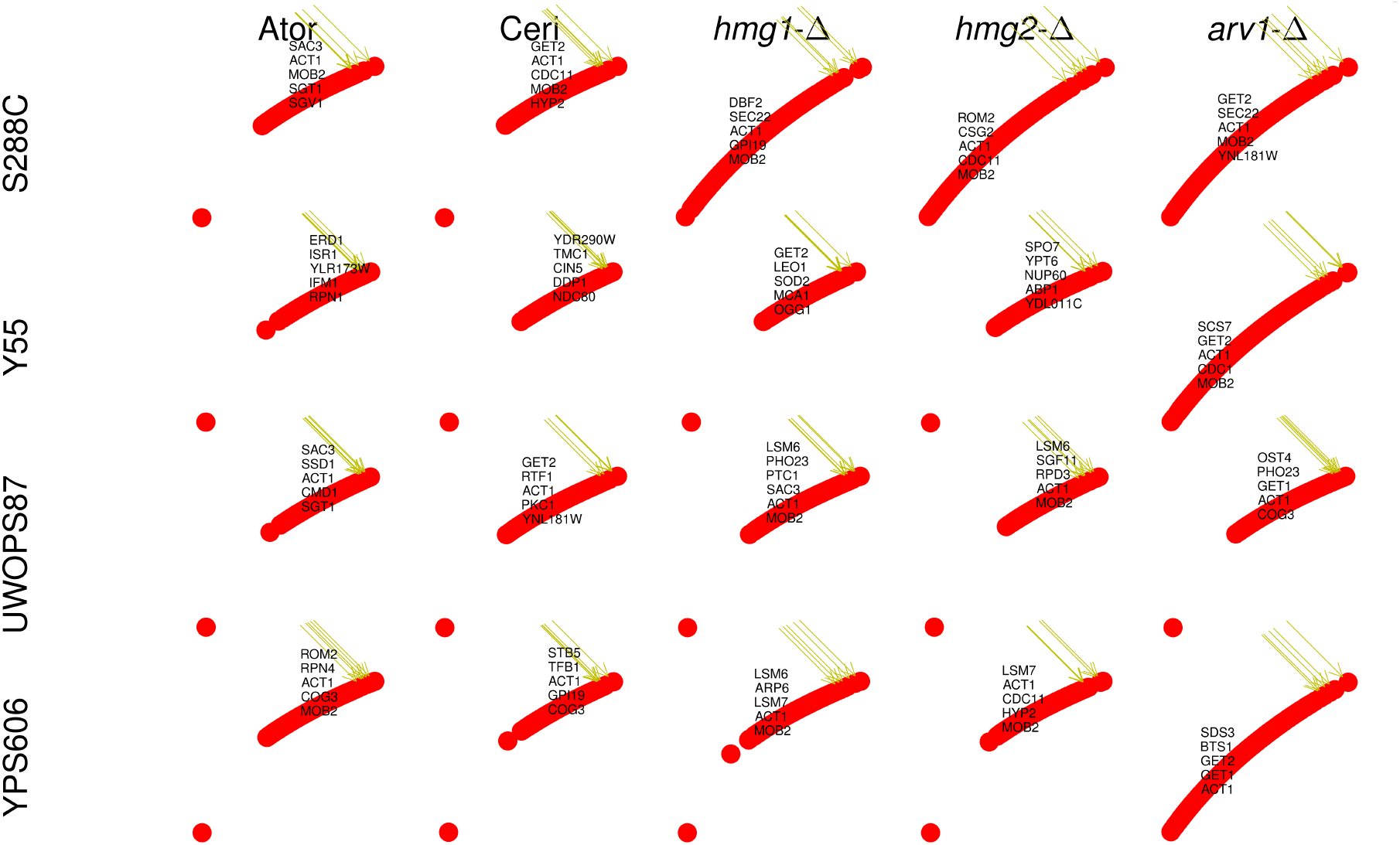
Deconvoluted master regulator genes with maximum closeness centrality for DNA libraries in four strains (S288C, Y55, UWOPS87-2421 (UWOPS87) and YPS606) by five queries (atorvastatin, cerivastatin, *hmg1-Δ, hmg2-Δ and arv1-Δ)*.

**Supplementary Figure S16.**
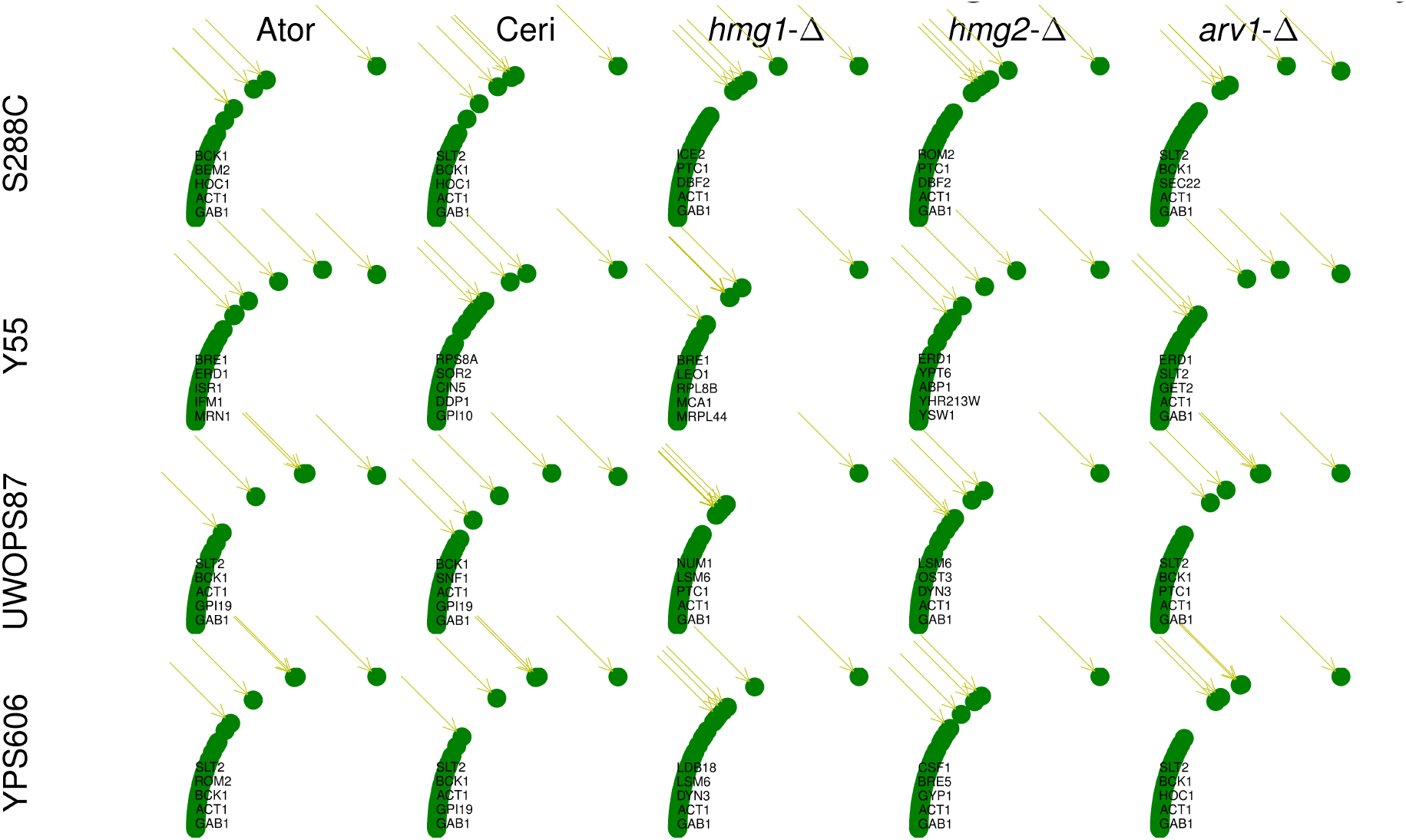
Deconvoluted master regulator genes with maximum eigenvector centrality for DNA libraries in four strains (S288C, Y55, UWOPS87-2421 (UWOPS87) and YPS606) by five queries (atorvastatin, cerivastatin, *hmg1-Δ*, *hmg2-Δ* and *arv1-Δ*).

**Supplementary Figure S17.**
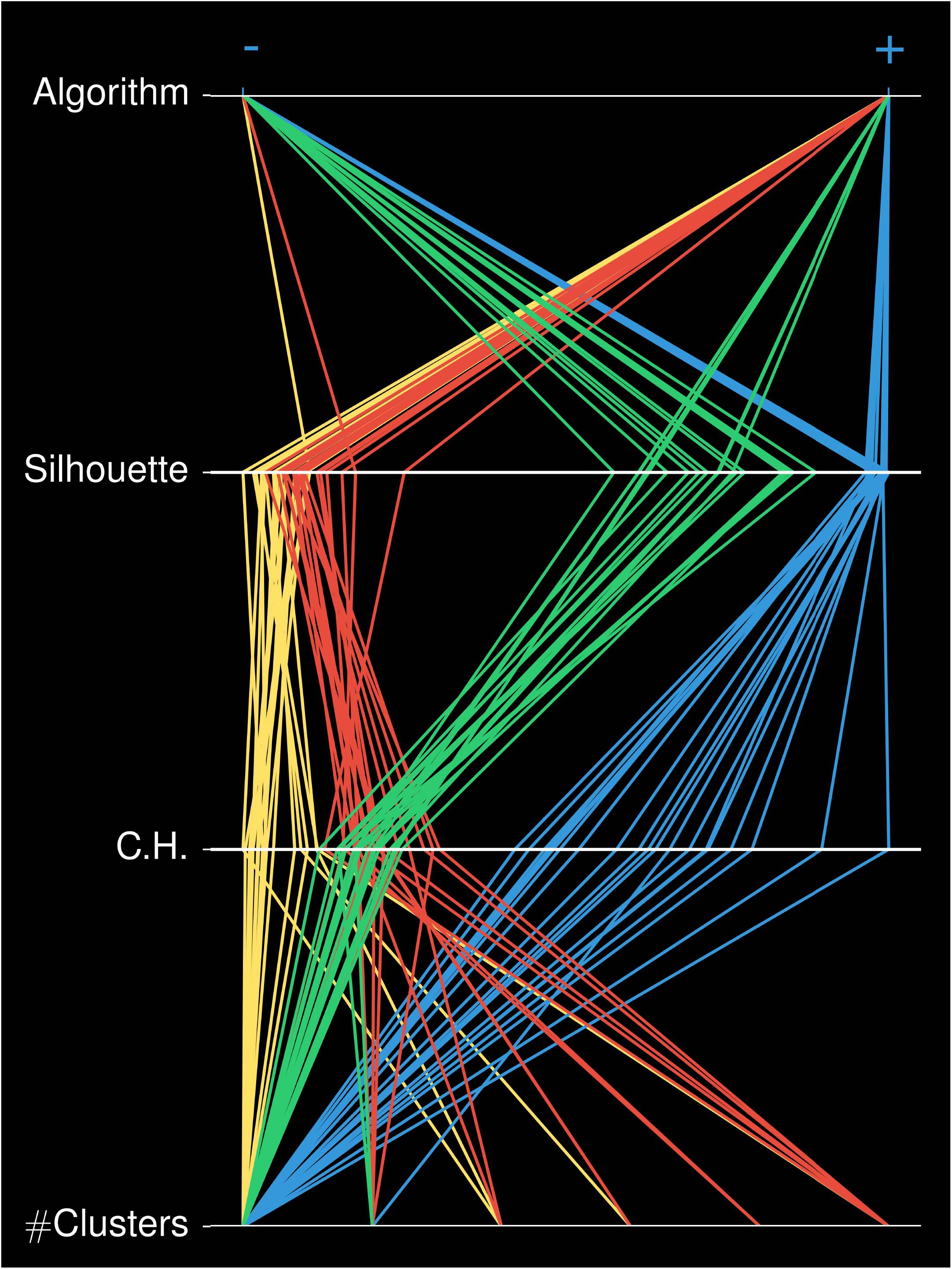
Genes from the primary GINs (Supplemental Table S9) were optimally clustered by topology values based on Silhouette score and C-H index into Cluster α (green), Cluster β (red), Cluster γ (yellow) and Cluster δ (blue). Note that cluster number variable starts at 3 on the LHS and ends at 8 on the RHS.

**Supplementary Figure S18.**
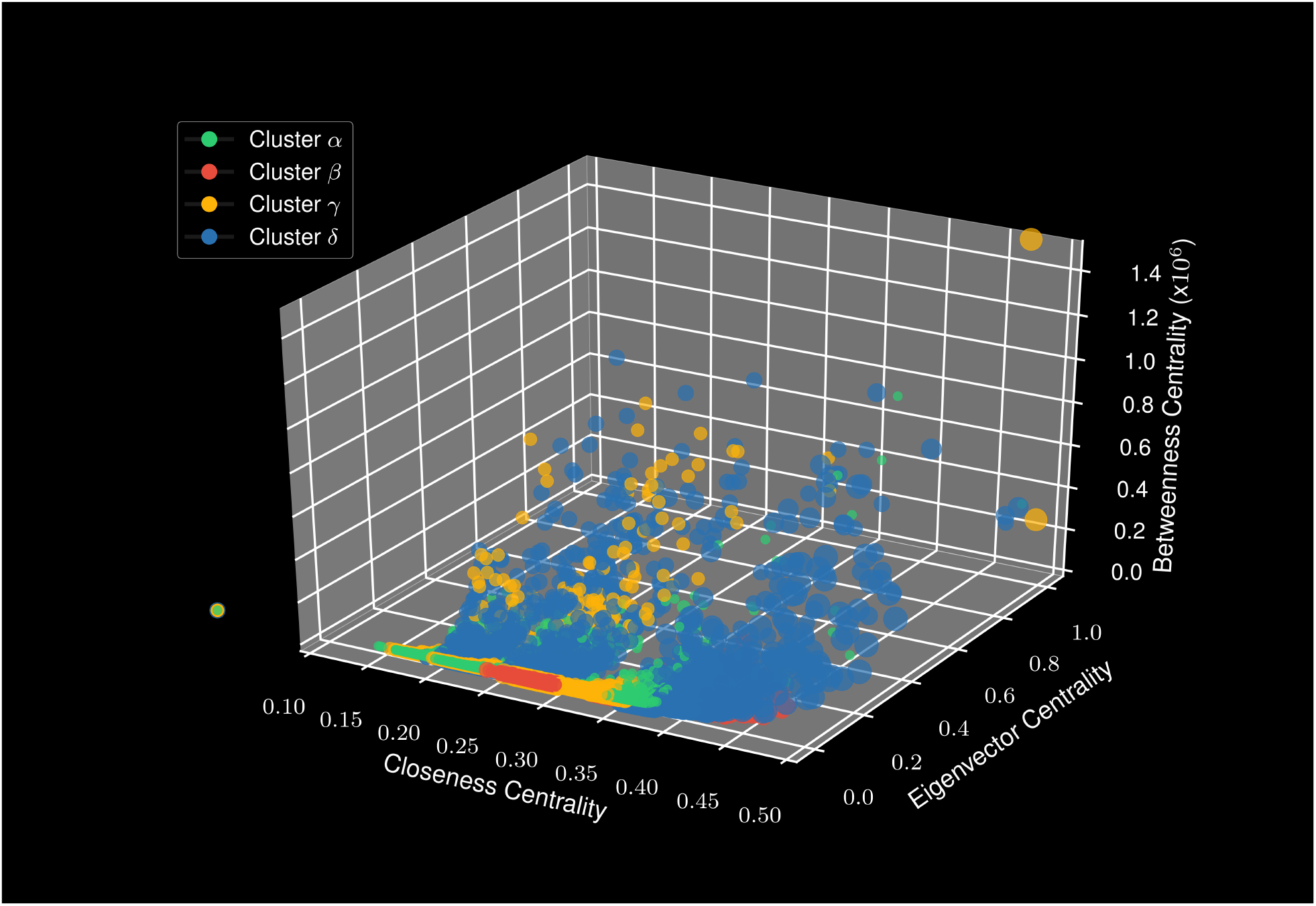
The four assigned clusters α, β, ϒ and δ were displayed on a 3-D plot by closeness, eigenvector and betweenness centralities, revealing maximum dispersion sensitivity to betweenness centrality values.

**Supplementary Figure S19.**
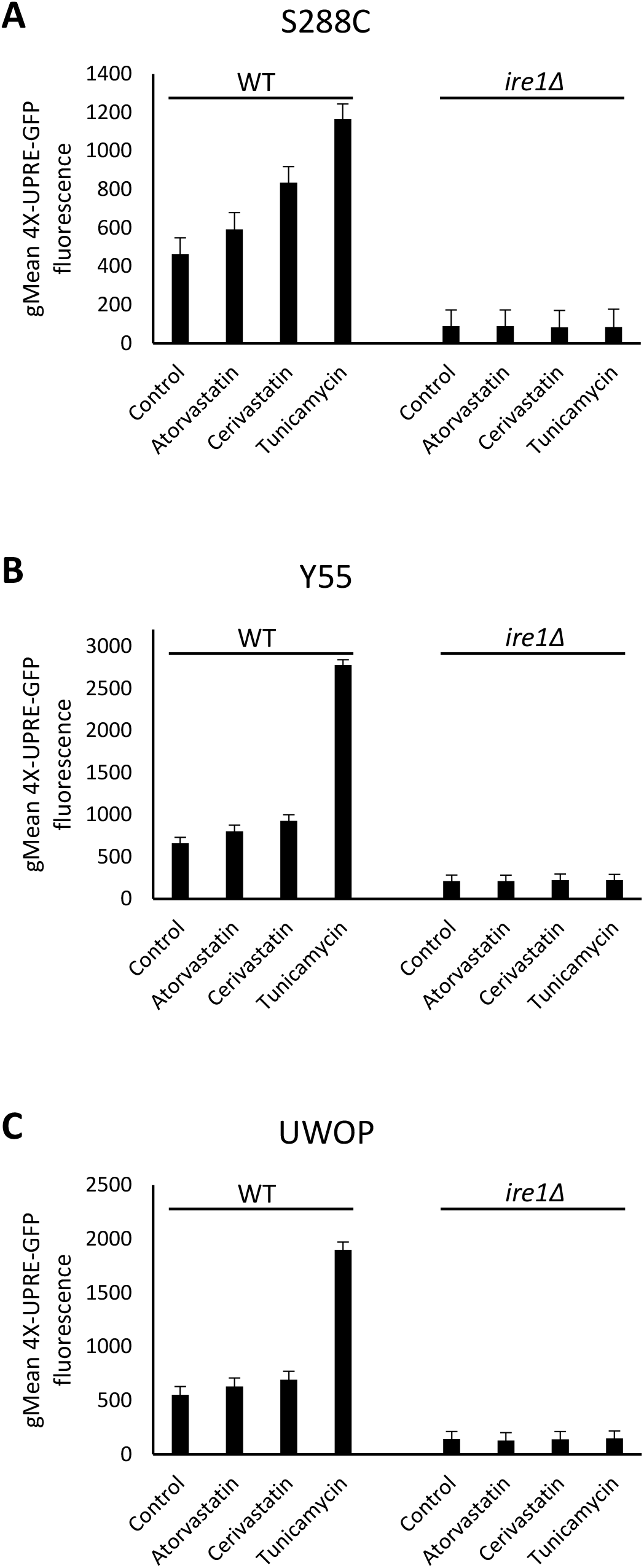
Flow cytometry of mid-log cells for the wild-type and *ire1-∆* strains of S288C, Y55 and UWOPS87-2421 (YPS606 was excluded due to increased flocculation impairing image analysis) expressing 4XUPRE-GFP cells and treated for 4 hours with atorvastatin, cerivastatin, or the established UPR inducer tunicamycin. The geometric mean (gMean) of each sample is shown with error bars representing the median absolute deviation.

## REFERENCES

1. Falconer, D.. & MacKay, T. F.. Introduction to Quantitative Genetics. (Longmans Green, Harlow, Essex, UK, 1996).

2. Yang, J. et al. Common SNPs explain a large proportion of the heritability for human height. Nat Genet 42, 565–569 (2010).

3. Chari, S. & Dworkin, I. The Conditional Nature of Genetic Interactions: The Consequences of Wild-Type Backgrounds on Mutational Interactions in a Genome-Wide Modifier Screen. PLOS Genetics 9, e1003661 (2013).

4. Dowell, R. D. et al. Genotype to Phenotype: A Complex Problem. Science 328, 469 (2010).

5. Forsberg, S. K. G., Bloom, J. S., Sadhu, M. J., Kruglyak, L. & Carlborg, Ö. Accounting for genetic interactions improves modeling of individual quantitative trait phenotypes in yeast. Nat Genet 49, 497–503 (2017).

6. Zuk, O. et al. Searching for missing heritability: designing rare variant association studies. Proc. Natl. Acad. Sci. U.S.A. 111, E455–464 (2014).

7. Mackay, T. F. C. Epistasis and quantitative traits: using model organisms to study gene-gene interactions. Nat Rev Genet 15, 22–33 (2014).

8. Mackay, T. F. & Moore, J. H. Why epistasis is important for tackling complex human disease genetics. Genome Medicine 6, 42 (2014).

9. Bloom, J. S., Ehrenreich, I. M., Loo, W. T., Lite, T.-L. V. & Kruglyak, L. Finding the sources of missing heritability in a yeast cross. Nature 494, 234–237 (2013).

10. Boone, C., Bussey, H. & Andrews, B. J. Exploring genetic interactions and networks with yeast. Nat Rev Genet 8, 437–449 (2007).

11. Baryshnikova, A., Costanzo, M., Myers, C. L., Andrews, B. & Boone, C. Genetic interaction networks: toward an understanding of heritability. Annual review of genomics and human genetics 14, 111–33 (2013).

12. Baryshnikova, A. Systematic Functional Annotation and Visualization of Biological Networks. Cell Syst 2, 412–421 (2016).

13. Costanzo, M. et al. A global genetic interaction network maps a wiring diagram of cellular function. Science 353, aaf1420 (2016).

14. Santolini, M. & Barabási, A.-L. Predicting perturbation patterns from the topology of biological networks. PNAS 201720589 (2018). doi:10.1073/pnas.1720589115

15. van Pel, D. M. et al. An evolutionarily conserved synthetic lethal interaction network identifies FEN1 as a broad-spectrum target for anticancer therapeutic development. PLoS Genet. 9, e1003254 (2013).

16. Tischler, J., Lehner, B. & Fraser, A. G. Evolutionary plasticity of genetic interaction networks. Nat Genet 40, 390–391 (2008).

17. Dixon, S. J. et al. Significant conservation of synthetic lethal genetic interaction networks between distantly related eukaryotes. Proceedings of the National Academy of Sciences of the United States of America 105, 16653–16658 (2008).

18. Roguev, A. et al. Conservation and rewiring of functional modules revealed by an epistasis map in fission yeast. Science (New York, N.Y.) 322, 405–10 (2008).

19. Wenner Moyer, M. The search beyond statins. Nat Med 16, 150–153 (2010).

20. Baker, S. K. Molecular clues into the pathogenesis of statin-mediated muscle toxicity. Muscle & nerve 31, 572–80 (2005).

21. Furberg, C. & Pitt, B. Withdrawal of cerivastatin from the world market. Current Controlled Trials in Cardiovascular Medicine 2, 205–207 (2001).

22. Karlson, B. W. et al. Variability of low-density lipoprotein cholesterol response with different doses of atorvastatin, rosuvastatin, and simvastatin: results from VOYAGER. Eur Heart J Cardiovasc Pharmacother 2, 212–217 (2016).

23. Hampton, R. Y. & Bhakta, H. Ubiquitin-mediated regulation of 3-hydroxy-3-methylglutaryl-CoA reductase. Proceedings of the National Academy of Sciences 94, 12944–12948 (1997).

24. Hampton, R. Y. & Rine, J. Regulated degradation of HMG-CoA reductase, an integral membrane protein of the endoplasmic reticulum, in yeast. J. Cell Biol. 125, 299–312 (1994).

25. Maciejak, A. et al. The effects of statins on the mevalonic acid pathway in recombinant yeast strains expressing human HMG-CoA reductase. BMC Biotechnology 13, 68 (2013).

26. Lee, M. N. et al. Common genetic variants modulate pathogen-sensing responses in human dendritic cells. Science 343, 1246980 (2014).

27. Parsons, A. B. et al. Exploring the Mode-of-Action of Bioactive Compounds by Chemical-Genetic Profiling in Yeast. Cell 126, 611–625 (2006).

28. Tong, A. H. Y. et al. Systematic Genetic Analysis with Ordered Arrays of Yeast Deletion Mutants. Science 294, 2364–2368 (2001).

29. Cubillos, F. A., Louis, E. J. & Liti, G. Generation of a large set of genetically tractable haploid and diploid Saccharomyces strains. 9, (2009).

30. Consortium, T. 1000 G. P. A map of human genome variation from population-scale sequencing. Nature 467, 1061–1073 (2010).

31. Schreiber, S. L. Chemical genetics resulting from a passion for synthetic organic chemistry. Bioorganic & Medicinal Chemistry 6, 1127–1152 (1998).

32. Cherry, J. M. et al. Saccharomyces Genome Database: the genomics resource of budding yeast. Nucleic acids research 40, D700–5 (2012).

33. D’Alessio, C., Caramelo, J. J. & Parodi, A. J. UDP-GlC:glycoprotein glucosyltransferase-glucosidase II, the ying-yang of the ER quality control. Semin. Cell Dev. Biol. 21, 491–499 (2010).

34. Walter, P. & Ron, D. The unfolded protein response: from stress pathway to homeostatic regulation. Science 334, 1081–1086 (2011).

35. Behnke, J., Feige, M. J. & Hendershot, L. M. BiP and its nucleotide exchange factors Grp170 and Sil1: mechanisms of action and biological functions. J. Mol. Biol. 427, 1589–1608 (2015).

36. Bircham, P. W. et al. Secretory pathway genes assessed by high-throughput microscopy and synthetic genetic array analysis. Molecular bioSystems 7, 2589–98 (2011).

37. Jonikas, M. C. et al. Comprehensive Characterization of Genes Required for Protein Folding in the Endoplasmic Reticulum. Science 323, 1693–1697 (2009).

38. Shechtman, C. F. et al. Loss of subcellular lipid transport due to ARV1 deficiency disrupts organelle homeostasis and activates the unfolded protein response. The Journal of biological chemistry 286, 11951–9 (2011).

39. Cronin, S. R., Rao, R. & Hampton, R. Y. Cod1p/Spf1p is a P-type ATPase involved in ER function and Ca2+ homeostasis. J. Cell Biol. 157, 1017–1028 (2002).

40. Kamada, T. & Kawai, S. An algorithm for drawing general undirected graphs. Information Processing Letters 31, 7–15 (1989).

41. Murtagh, F. & Legendre, P. Ward’s Hierarchical Agglomerative Clustering Method: Which Algorithms Implement Ward’s Criterion? J Classif 31, 274–295 (2014).

42. Hartwell, L. H., Hopfield, J. J., Leibler, S. & Murray, A. W. From molecular to modular cell biology. Nature 402, C47–52 (1999).

43. Proulx, S. R., Promislow, D. E. L. & Phillips, P. C. Network thinking in ecology and evolution. Trends Ecol. Evol. (Amst.) 20, 345–353 (2005).

44. Sah, P., Singh, L. O., Clauset, A. & Bansal, S. Exploring community structure in biological networks with random graphs. BMC Bioinformatics 15, 220 (2014).

45. De Meo, P., Ferrara, E., Fiumara, G. & Ricciardello, A. A novel measure of edge centrality in social networks. Knowledge-Based Systems 30, 136–150 (2012).

46. Barabási, A.-L., Gulbahce, N. & Loscalzo, J. Network medicine: a network-based approach to human disease. Nature Reviews Genetics 12, 56–68 (2011).

47. Özgür, A., Vu, T., Erkan, G. & Radev, D. R. Identifying gene-disease associations using centrality on a literature mined gene-interaction network. Bioinformatics 24, i277–i285 (2008).

48. Mclean, C., He, X., T, I. S. & J, D. A. Improved Functional Enrichment Analysis of Biological Networks using Scalable Modularity Based Clustering. Journal of Proteomics & Bioinformatics 9, (2016).

49. Rousseeuw, P. J. Silhouettes: A graphical aid to the interpretation and validation of cluster analysis. Journal of Computational and Applied Mathematics 20, 53–65 (1987).

50. Supek, F., Bošnjak, M., Škunca, N. & Šmuc, T. REVIGO Summarizes and Visualizes Long Lists of Gene Ontology Terms. PLOS ONE 6, e21800 (2011).

51. Kolaczkowski, M., Kolaczowska, A., Luczynski, J., Witek, S. & Goffeau, A. In vivo characterization of the drug resistance profile of the major ABC transporters and other components of the yeast pleiotropic drug resistance network. Microb. Drug Resist. 4, 143–158 (1998).

52. Travers, K. J. et al. Functional and genomic analyses reveal an essential coordination between the unfolded protein response and ER-associated degradation. Cell 101, 249–258 (2000).

53. Deutschbauer, A. M. & Davis, R. W. Quantitative trait loci mapped to single-nucleotide resolution in yeast. Nat Genet 37, 1333–1340 (2005).

54. Cubillos, F. A. et al. High-Resolution Mapping of Complex Traits with a Four-Parent Advanced Intercross Yeast Population. Genetics 195, 1141–1155 (2013).

55. Roberts, C. A., Miller, J. H. & Atkinson, P. H. The genetic architecture in Saccharomyces cerevisiae that contributes to variation in drug response to the antifungals benomyl and ketoconazole. FEMS Yeast Res. 17, (2017).

56. Bandyopadhyay, S. et al. Rewiring of genetic networks in response to DNA damage. Science 330, 1385–1389 (2010).

57. van Opijnen, T., Dedrick, S. & Bento, J. Strain Dependent Genetic Networks for Antibiotic-Sensitivity in a Bacterial Pathogen with a Large Pan-Genome. PLoS Pathog. 12, e1005869 (2016).

58. Promlek, T. et al. Membrane aberrancy and unfolded proteins activate the endoplasmic reticulum stress sensor Ire1 in different ways. Mol. Biol. Cell 22, 3520–3532 (2011).

59. Kanemoto, S. et al. Multivesicular body formation enhancement and exosome release during endoplasmic reticulum stress. Biochem. Biophys. Res. Commun. 480, 166–172 (2016).

60. Araki, M., Maeda, M. & Motojima, K. Hydrophobic statins induce autophagy and cell death in human rhabdomyosarcoma cells by depleting geranylgeranyl diphosphate. Eur. J. Pharmacol. 674, 95–103 (2012).

61. Boucher, B. & Jenna, S. Genetic interaction networks: better understand to better predict. Frontiers in Genetics 4, (2013).

62. Boucher, B., Lee, A. Y., Hallett, M. & Jenna, S. Structural and Functional Characterization of a Caenorhabditis elegans Genetic Interaction Network within Pathways. PLoS Comput. Biol. 12, e1004738 (2016).

63. Rizzolo, K. et al. Systems analysis of the genetic interaction network of yeast molecular chaperones. Mol. Omics 14, 82–94 (2018).

64. Wood, A. R. et al. Defining the role of common variation in the genomic and biological architecture of adult human height. Nat. Genet. 46, 1173–1186 (2014).

65. Schoenrock, A. et al. Evolution of protein-protein interaction networks in yeast. PLoS ONE 12, e0171920 (2017).

66. Bridgham, J. T., Ortlund, E. A. & Thornton, J. W. An epistatic ratchet constrains the direction of glucocorticoid receptor evolution. Nature 461, 515–519 (2009).

67. Amberg, D. C., Burke, D. & Strathern, J. N. Methods in Yeast Genetics: A Cold Spring Harbor Laboratory Course Manual. (CSHL Press, 2005).

68. Goldstein, A. L. & McCusker, J. H. Three new dominant drug resistance cassettes for gene disruption in Saccharomyces cerevisiae. Yeast 15, 1541–1553 (1999).

69. Engel, S. R. et al. The reference genome sequence of Saccharomyces cerevisiae: then and now. G3 (Bethesda) 4, 389–398 (2014).

70. Liti, G. et al. Population genomics of domestic and wild yeasts. Nature 458, 337–341 (2009).

71. Poliakov, A., Foong, J., Brudno, M. & Dubchak, I. GenomeVISTA--an integrated software package for whole-genome alignment and visualization. Bioinformatics 30, 2654–2655 (2014).

72. Wagih, O. & Parts, L. gitter: a robust and accurate method for quantification of colony sizes from plate images. G3 (Bethesda, Md.) 4, 547–52 (2014).

73. Dittmar, J. C., Reid, R. J. & Rothstein, R. ScreenMill: a freely available software suite for growth measurement, analysis and visualization of high-throughput screen data. BMC bioinformatics 11, 353 (2010).

74. Shannon, P. et al. Cytoscape: a software environment for integrated models of biomolecular interaction networks. Genome Res. 13, 2498–2504 (2003).

75. Kuzmin, E. et al. Systematic analysis of complex genetic interactions. Science 360, (2018).

76. Blondel, V. D., Guillaume, J.-L., Lambiotte, R. & Lefebvre, E. Fast unfolding of communities in large networks. J. Stat. Mech. 2008, P10008 (2008).

77. Klopfenstein, D. V. et al. GOATOOLS: A Python library for Gene Ontology analyses. Scientific Reports 8, 10872 (2018).

78. Lin, D. An Information-Theoretic Definition of Similarity. in Proceedings of the Fifteenth International Conference on Machine Learning 296–304 (Morgan Kaufmann Publishers Inc., 1998).

79. Cox, J. S. & Walter, P. A Novel Mechanism for Regulating Activity of a Transcription Factor That Controls the Unfolded Protein Response. Cell 87, 391–404 (1996).

80. Bicknell, A. A., Babour, A., Federovitch, C. M. & Niwa, M. A novel role in cytokinesis reveals a housekeeping function for the unfolded protein response. J. Cell Biol. 177, 1017–1027 (2007).

81. Yang, J. et al. Common SNPs explain a large proportion of the heritability for human height. Nat Genet 42, 565–569 (2010).

